# Differential GLP-1R binding and activation by peptide and non-peptide agonists

**DOI:** 10.1101/2020.08.16.252585

**Authors:** Xin Zhang, Matthew J. Belousoff, Peishen Zhao, Albert J. Kooistra, Tin T. Truong, Sheng Yu Ang, Christina Rye Underwood, Thomas Egebjerg, Petr Šenel, Gregory D. Stewart, Yi-Lynn Liang, Alisa Glukhova, Hari Venugopal, Arthur Christopoulos, Sebastian G. B. Furness, Laurence J. Miller, Steffen Reedtz-Runge, Christopher J. Langmead, David E. Gloriam, Radostin Danev, Patrick M. Sexton, Denise Wootten

## Abstract

Peptide drugs targeting class B1 GPCRs can treat multiple diseases, however there remains substantial interest in the development of orally delivered non-peptide drugs. Here we reveal unexpected overlap between signalling and regulation of the glucagon-like peptide-1 (GLP-1) receptor by the non-peptide agonist, PF 06882961, and GLP-1 that was not observed for another compound, OWL-833. Both compounds are currently in clinical trials for treatment of type 2 diabetes. High resolution cryo-EM structures reveal the binding sites for PF-06882961 and GLP-1 substantially overlap, whereas OWL-833 adopts a unique binding mode with a more open receptor conformation at the extracellular face. Structural differences involving extensive water-mediated hydrogen bond networks could be correlated to functional data to understand how PF 06882961, but not OWL-833, can closely mimic the pharmacological properties of GLP-1. These findings will facilitate rational structure-based discovery of non-peptide agonists targeting class B GPCRs.

## INTRODUCTION

Class B1 G protein-coupled receptors (GPCRs) are activated by peptide hormones/neuropeptides and regulate physiological functions including glycaemic control, satiety, neuroprotection, bone turnover and cardiovascular tone (Culhane et al., 2015; Pal et al., 2012). These receptors predominantly couple to the stimulatory protein G_s_ to promote cAMP production, however they can activate a range of signalling pathways and are highly susceptible to biased agonism (Wootten et al., 2017). In the past three years numerous active state structures of class B1 GPCRs coupled to the stimulatory protein G_s_ have been determined using single particle cryo-electron microscopy (cryo-EM) (Liang et al., 2020a; Liang et al., 2020b; Liang et al., 2018a; Liang et al., 2018b; Liang et al., 2017; Ma et al., 2020; Qiao et al., 2020; Zhang et al., 2017; Zhao et al., 2019). The resulting structures revealed common peptide binding pockets and a conserved G_s_ binding site within the receptor transmembrane (TM) helix bundle. However, distinct conformations have been identified within the extracellular face of different receptors that may be linked to evolutionary divergence and receptor selectivity (Liang et al., 2020b).

The glucagon peptide-1 receptor (GLP-1R) is a class B1 GPCR that is the target of multiple clinically approved peptide drugs for the treatment of type 2 diabetes, with select drugs also approved for obesity (Brown et al., 2018; Nauck and Meier, 2019). These peptides have been engineered to enhance pharmacokinetic profiles over the rapidly cleared native GLP-1 peptides, while maintaining potency at the GLP-1R. Nonetheless, there are differences in clinical efficacies of these drugs and some are also able to reduce body weight in obese patients, improve cardiovascular outcomes, and may have benefit for the treatment of neurodegenerative disorders (Brown et al., 2018; Grieco et al., 2019; Nauck and Meier, 2019). To date, the structures of active state GLP-1R:G_s_ complexes bound to either the native GLP-1 peptide or a G protein biased agonist, exendin-P5 (ExP5) have been determined using cryo-EM (Liang et al., 2018b; Zhang et al., 2017). In addition, a crystal structure of the inactive full-length GLP-1R bound by an extracellular antibody, in the absence of a peptide ligand, was recently reported (Wu et al., 2020). These structures provide critical information regarding GLP-1R activation by peptide agonists and combined with molecular dynamics simulations and molecular pharmacology have provided initial insights into the molecular basis of differential efficacy and biased agonism induced by peptide ligands.

With the exception of semaglutide, which is also approved in oral formulation, all GLP-1 peptide analogues require parenteral administration. Moreover, these peptides are expensive to produce and induce nausea and other gastrointestinal problems that reduce patient compliance (Pearson et al., 2019). Consequently, the identification and development of orally available small molecules that can also broadly mimic the actions of GLP-1 is of major interest. Recently, several small molecule GLP-1R agonists have entered into clinical trials, including PF 06882961 (Pfizer), TTP-273 (vTv therapeutics/Huadong medicine Co Ltd) and OWL-833 (Chugai/Eli Lilly). We revealed that TT-OAD2, an analogue reported in the patent series that includes TTP-273, displays unique kinetic and signalling properties, and has a distinct binding mode relative to GLP-1 (Zhao et al., 2020). This demonstrated that ligand engagement deep within the TM core was not a requirement for class B1 GPCR activation from the extracellular face. Currently, very little is known regarding the binding mode, activation and signalling profiles of PF 06882961 and OWL-833. These molecules have greater activity at the GLP-1R than TT-OAD2 and therefore may hold more promise in the clinic. Here we revealed that PF 06882961 has a remarkably similar overall signalling and regulatory profile to GLP-1, whereas OWL-833 exhibited a distinct profile that is strongly biased towards G_s_-mediated cAMP production relative to the spectrum of responses elicited by GLP-1. High resolution structural information on the active GLP-1R-G_s_ signalling complex in the presence of GLP-1, PF 06882961 and OWL-833 provide unique mechanistic insights into the divergent binding modes of each of these agonists and can explain how PF 06882961, but not OWL-833, is able to mimic the actions of GLP-1.

## RESULTS AND DISCUSSION

### PF 06882961 and OWL-833 have different biased agonism profiles and kinetics of GLP-1R activation

The GLP-1R is pleiotropically coupled to multiple transducer and regulatory proteins, including G proteins, receptor kinases and *β*-arrestins (Graaf et al., 2016). Despite their advancement in clinical trials, the pharmacodynamic properties of PF 06882961 and OWL-833 have not been reported and the extent to which these compounds overlap the signalling and regulatory profile of endogenous GLP-1 is unknown. To address this, using GLP-1 as a reference, we performed time-resolved assays of well-studied downstream signalling and regulatory events linked to GLP-1R activation; cAMP production, pERK1/2, intracellular calcium mobilisation, *β*-arrestin-1 and *β*-arrestin-2 recruitment. We also monitored ligand-induced G_s_ conformational changes (associated with G_s_ activation and cAMP signalling) and receptor trafficking into FYVE-containing early endosomes, where continued peptide-mediated signalling can contribute to cellular response (**Figures 1 & S1**).

**Figure 1.**
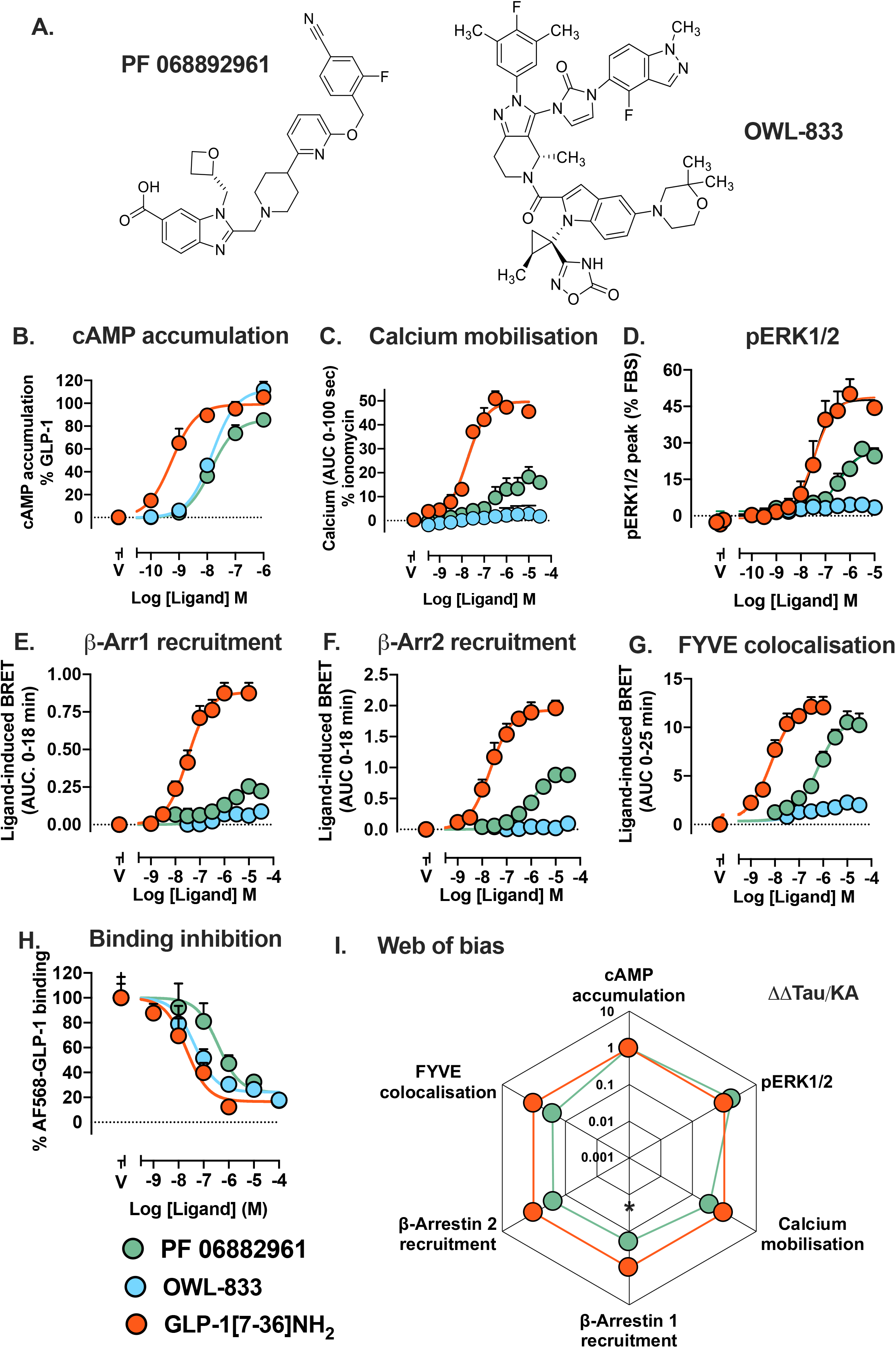
Pharmacological profiles of GLP-1, PF 06882961 and OWL-833. **A,** Chemical structures of PF 06882961 and OWL-833. **B-G**, Concentration response curves showing the signalling profiles of GLP-1 and non-peptidic agonists in cAMP accumulation (B), intracellular calcium mobilisation (C), pERK1/2 (D), *β*-arrestin-1 recruitment (E), *β*-arrestin-2 recruitment (F) and GLP-1R internalisation (as measured by ligand-induced colocalization with the early endosome marker, FYVE) (G). Panels C-G were plotted using the area under the curve (AUC) analysis applied to data from Figure S1. **H,** Inhibition binding curves showing the ability of PF 06882961 and OWL-833 to inhibit the BRET signal between NLuc-GLP-1R and AF568-GLP-1. Data were derived from the last time point in Figure S1G. **I**, “web of bias” plots *ΔΔτ*/K_A_ values on a logarithmic scale showing the signalling in different pathways of PF-06882961 relative to the reference agonist GLP-1 and the reference pathway cAMP. *τ*/K_A_ were calculated by applying the operational model of agonism to data in panels B-G. * indicates statistically significant bias (P < 0.05) relative to GLP-1 as determined by a two-way ANOVA followed by a Sidak’s multiple comparisons test, comparing each pathway to cAMP and PF 06882961 to GLP-1. All data are the means + S.E.M of five independent experiments performed in duplicate or triplicate (with the exception of panel H, where 3 independent experiments were performed). *See also Figure S1*.

In a HEK293 stable cell line overexpressing the GLP-1R, PF 06882961 and OWL-833 exhibited equivalent potency and were full agonists for cAMP production, with ∼30-fold lower potency than GLP-1 (**Figure 1B**). This was correlated with similar potencies for the two non-peptides in inducing conformational changes in G_s_ (**Figure S1A**), however, OWL-833 was inactive in all other assays at concentrations up to 10 μM (**Figure 1C-G & S1**). In marked contrast, PF 06882961 was active in all assays (30-100-fold lower potency than GLP-1), displaying similar maximal internalisation to GLP-1, but only partial agonism in pERK1/2, calcium mobilisation and *β*-arrestin recruitment (**Figures 1B-1G & S1**). Quantification of responses using operational modelling revealed subtle biased agonism of PF 06882961, relative to GLP-1 that was only significantly different for *β*-arrestin-1 recruitment relative to cAMP (**Figure 1I**). However, examination of the kinetics of observed responses revealed that while PF 06882961 had a similar profile to GLP-1 for induction of G protein conformational changes, calcium signalling and pERK1/2, it was slower at inducing *β*-arrestin recruitment and internalisation (**Figure S1**). Nonetheless, given the major divergence in the chemical properties of the small molecule agonist (**Figure 1A**), and the GLP-1 peptide, overall PF 06882961 exhibited a remarkably similar signalling and regulatory profile to GLP-1 (**Figure 1I**). Indeed, many peptide agonists, including the clinically approved peptides, exendin-4 and liraglutide, and the low potency natural agonist, oxyntomodulin, demonstrate greater biased agonism across the assessed pathways than PF 06882961 compared to GLP-1 (Fletcher et al., 2018; Hager et al., 2016; Koole et al., 2010).

Due to the lack of response in the majority of pathways, bias could not be quantified for OWL-833, but, as it had an equivalent response to PF 06882961 in cAMP production, we can infer that OWL-833 is heavily biased towards this pathway, and away from all other pathways assessed, when compared to GLP-1 and PF 06882961. Interestingly, assessment of binding properties in a hemi-equilibrium membrane binding assay revealed that OWL-833 had ∼10-fold higher affinity than PF 06882961 (**Figures 1H & S1G**), suggesting that while the two agonists have similar concentration response profiles in cAMP accumulation (**Figure 1B**), PF 06882961 has higher efficacy for this pathway.

Intriguingly, OWL-833 has a similar signalling and regulatory profile to that of TT-OAD2, where neither agonist promoted *β*-arrestin recruitment or internalisation and both were also very poor agonists for pERK1/2 and calcium mobilisation. Nonetheless, OWL-833 was a more efficient agonist, displaying higher potency and maximal responses than TT-OAD2, for cAMP production (**Figure 1**) (Zhao et al., 2020). This may, at least in part, be due to the much slower rate of TT-OAD2-induced G_s_ conformational transitions linked to cAMP production, compared to OWL-833 (**Figure S1)** (Zhao et al., 2020).

Collectively, these data suggest that PF 06882961 is likely to have relatively similar pharmacodynamics properties to GLP-1 *in vivo*, while OWL-833 and TT-OAD2 may exhibit greater similarity in their pharmacodynamic profiles (albeit with markedly different kinetics for canonical G_s_-mediated signalling). However, these data also raise critical mechanistic and structural questions on how the different small molecule agonist engage and activate the GLP-1R, and in particular with respect to how PF 06882961 can mimic key properties of the native peptide.

### Determination of high resolution structures of distinct agonist-bound GLP-1R:G_s_ complexes

To understand how the different agonists interact with the GLP-1R, we determined human GLP-1R:G_s_ structures bound to GLP-1, PF 06882961 or OWL-833 (**Figures 2, S2**). Complexes were induced by the addition of high concentrations of agonist (10μM GLP-1 or 50μM for small molecules) prior to solubilisation and purification. Purified complexes were resolved as monodispersed peaks on size exclusion chromatography (SEC) and contained all of the expected components (**Figure S2**). Vitrified complexes were imaged by cryo-EM yielding 2D class averages with well resolved secondary structure that were reconstructed into 3D consensus density maps with global resolutions of 2.1 Å for GLP-1, 2.5 Å for PF 06882961 and 2.1 Å for OWL-833 at gold standard FSC 0.143 (**Figure S2, Table S1**). The GLP-1 bound complex provides a 2.0 Å increase in resolution over the previously published 4.1 Å map of the active GLP-1:rabbit GLP-1R:G_s_ complex that had limited density for much of the extracellular side of the receptor (Zhang et al., 2017). The *α*-helical domain (AHD) of the G*α*_s_ subunit was poorly resolved in all consensus maps and was masked out during the final refinement. To improve the resolution of domains that had higher mobility, and thus lower local resolution in consensus maps, localised 3D-refinements focused on the full receptor, the receptor extracellular domain (ECD), or the G protein were also performed (**Figure S2**). These focused maps enabled robust molecular modelling within the receptor ligand binding pockets and the ECD for each complex.

**Figure 2.**
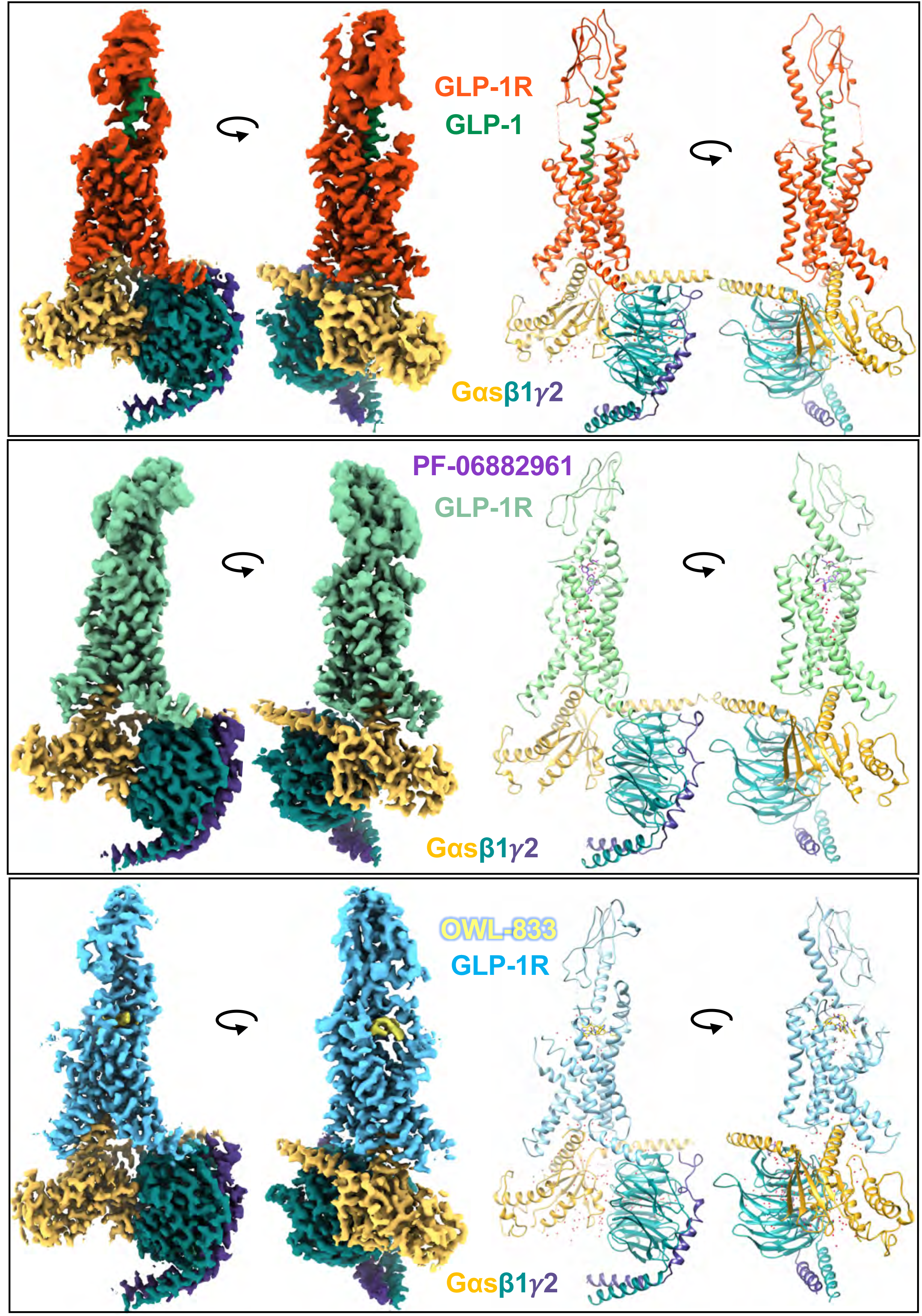
Cryo-EM structures of GLP-1R:G_s_ complexes. **Top**, GLP-1-bound; **Middle**, PF 06882961-bound; **Bottom**, OWL-833-bound. The left-hand side shows orthogonal views of the cryo-EM maps, while the right-hand side displays the backbone models built into the maps in ribbon format. Colouring denotes the protein segments as highlighted on the figure panels. *See also Figures 3, S2, S3 and S7 and Videos S1, S3 and S4*.

Using a combination of the consensus, full receptor refined, and ECD refined maps of the GLP-1-bound complex, atomic level modelling of the entire peptide, almost the full receptor and G protein, excepting the G*α*_s_ AHD was performed (**Figures S2-S4**). The GLP-1R intracellular loop (ICL) 3 (N338-D344) was poorly resolved in all maps, indicative of high mobility and was not modelled. Within the PF 06882961 and OWL-833 bound GLP-1R cryo-EM maps the GLP-1R TM domain was well resolved allowing unambiguous modelling of both ligands, and the majority of receptor residues, including the binding sites and the majority of the loop regions, with the exception of ICL3 in both complexes, where only the backbone was modelled, and the TM1/ECD linkers where S129-S134 was not modelled (**Figures 2 & S3**). ECL3 between H374 and T377 was also poorly resolved and not modelled for the OWL-833-bound receptor. The GLP-1R ECD, in both small molecule agonist complexes, was better resolved than in the GLP-1 bound map (**Figure S2**) and enabled unambiguous modelling of the backbone and the majority of side chain residues for both complexes using the receptor refined maps.

### The GLP-1 peptide forms extensive interactions with the receptor TM domain and structural waters

Structural understanding of the mechanism through which PF 06882961, but not OWL-833, can mimic the pharmacology of GLP-1 first requires molecular accuracy in understanding of how GLP-1 itself binds to the receptor, including the involvement of structural waters in mediating interactions. While the previously published GLP-1:GLP-1R:G_s_ structure, 5VAI (Zhang et al., 2017), is generally similar to our structure, particularly towards the intracellular face, there are substantial differences in the models built into the cryo-EM maps for the peptide binding site, ECLs and the top of TM1, consistent with difficulty in modelling these regions where there was limited resolution in the 4.1 Å map used for 5VAI (**Figure S4A**). As such, details of GLP-1 binding from our high-resolution structure that are well supported by density in the cryo-EM map (**Figures S3, S4**) are described below.

The N-terminal amino acids, H7^GLP-1^ and A8^GLP-1^ are crucial for GLP-1 function, as truncation of these two residues, forming the GLP-1(9-36)NH_2_ analogue, lowers affinity and efficacy (Montrose-Rafizadeh et al., 1997). These two residues reside deep within the TM cavity where they are located above the class B1 GPCR conserved central polar network, and form extensive interactions with GLP-1R residues (**Figure 3**). H7^GLP-1^ interacts with residues in TM3 and TM5 through hydrogen bond and hydrophobic interactions with Q234^3.37^, V237^3.40^, W306^5.36^, R310^5.40^ and I313^5.43^, whereas A8^GLP-1^ forms hydrophobic interactions with E387^7.42^ and L388^7.43^ in TM7 (**Figures 3 & S4**). In addition, H7^GLP-1^ is stabilised through a hydrogen bond of its backbone with the side chain of T11^GLP-1^ and through a structural water that interacts with the backbone of TM5. E9^GLP-1^ also resides at the base of the binding pocket participating in van der Waals interactions with R190^2.60^ and L388^7.43^ as well as hydrogen bond interactions with R190^2.60^ and Y152^1.47^, both crucial residues for GLP-1R activation and signalling (Lei et al., 2018; Wootten et al., 2016a; Wootten et al., 2016b). In addition, A8^GLP-1^ (backbone) and E9^GLP-1^ engage in an extensive hydrogen bond network with structural waters and polar residues at the base of the peptide binding pocket (R190^2.60^, Y152^1.47^, Y241^3.44^), as well as with the backbone of TM7 (**Figures 3 & S4**). This water mediated interaction network likely stabilises the peptide in the base of the pocket, and helps maintain an active receptor conformation, as alanine mutation of these receptor side chains reduced both affinity and efficacy of GLP-1 and other peptide ligands (**Figure S5**) (Lei et al., 2018; Wootten et al., 2016a; Wootten et al., 2016b). The GLP-1 peptide also forms extensive interactions with ECLs 2 and 3 where S14^GLP-1^ and S17^GLP-1^ interact via both hydrogen bonding and van der Waals interactions with T298^ECL2^, R299^ECL2^ and N300^ECL2^. T11^GLP-1^ and D15^GLP-1^ form polar and hydrophobic interactions with D372^ECL3^, R380^ECL3/7.35^ and L384^7.39^ (**Figures 3 & S4**). Residues at these positions within ECL2 and 3 are crucial for peptide affinity and receptor activation in numerous class B1 GPCRs (Coin et al., 2013; Koole et al., 2012a; Woolley et al., 2017). Other notable interactions include multiple hydrophobic contacts of the peptide with residues in TM1 (L141^1.36^, L144^1.39^, Y148^1.43^), hydrogen bonding between K197^2.67^ and T13^GLP-1^, and Y205^2.75^ with both S17^GLP-1^ and E21^GLP-1^. E21^GLP-1^ also interacts with the backbone of the far N-terminus of the receptor. The importance of these key interactions in GLP-1 function are supported by extensive alanine mutagenesis studies (**Figure S5**) (Coopman et al., 2010; Dods and Donnelly, 2015; Graaf et al., 2016; Lei et al., 2018; Wootten et al., 2016c; Wootten et al., 2013b; Yang et al., 2016).

**Figure 3.**
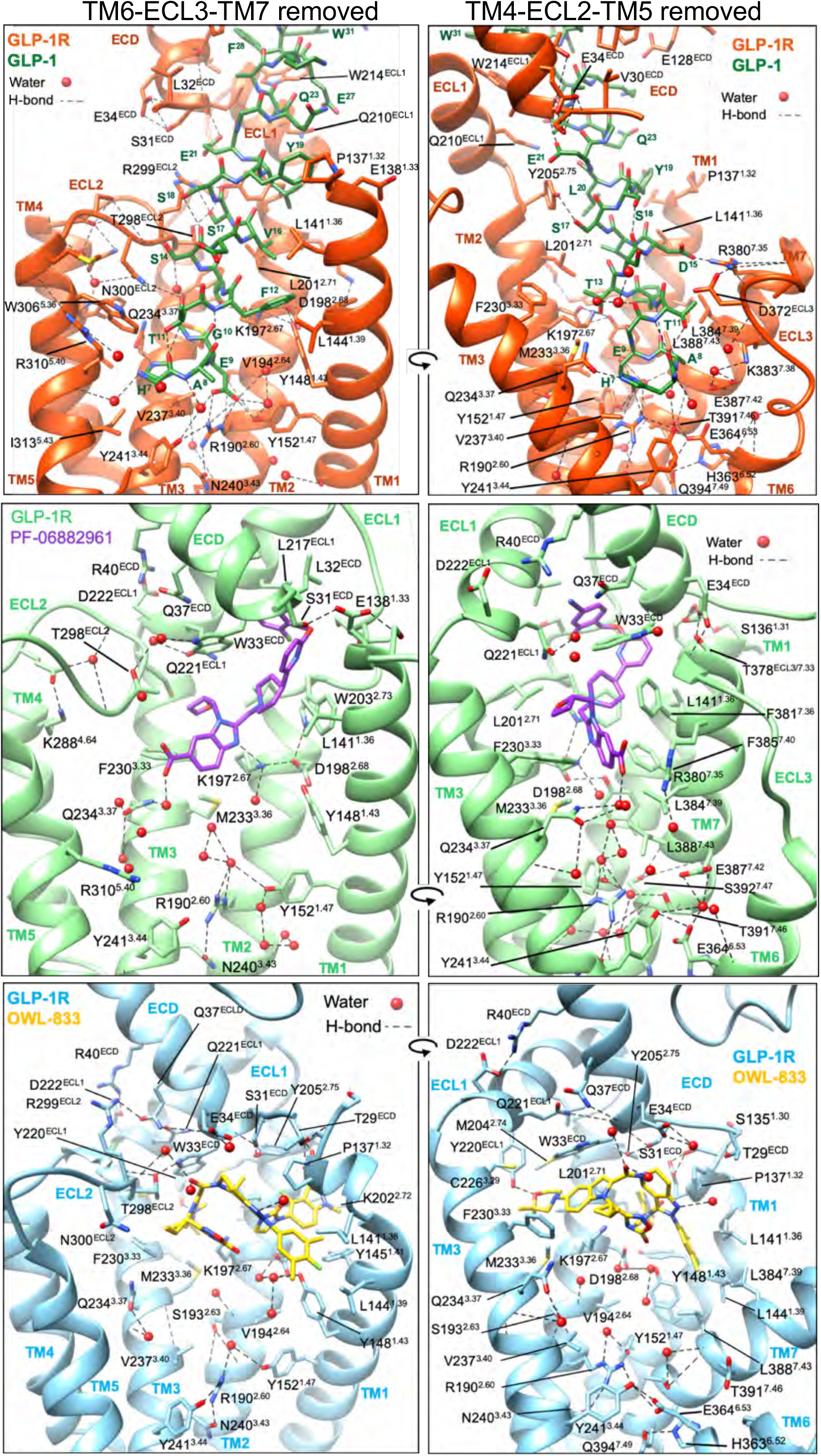
Interactions of GLP-1 and non-peptidic agonists within the TM binding cavity of the GLP-1R. **Top,** GLP-1R residues that interact with ligand or waters (red balls) within the GLP-1R binding cavity are displayed in stick format. **Top**, GLP-1R (orange) and GLP-1 peptide (green); **Middle,** GLP-1R (pale green) and PF 06882961 (purple); **Bottom,** GLP-1R (pale blue) and OWL-833 (yellow). For each binding site two views are depicted for clarity; Left; side view of the TM bundle viewed from the upper portion of TM6/TM7 where TM6-ECL3-TM7 have been removed; Right, side view of the TM bundle viewed from the upper portion of TM4/TM5 where TM4-ECL2-TM5 have been removed. Dashed lines depict hydrogen bonds as determined using UCSF chimera. *See also Figures 2, S3, S4 and S5*.

### PF 06882961 but not OWL-833 mimics key features of the GLP-1 binding linked to receptor activation and biased agonism

Relative to GLP-1, both small molecule agonists bind superficially to the transmembrane domain of the receptor but adopt very distinct poses where they engage in different receptor interactions (detailed in **Figures 3, S4 & Table S2**) and induce distinct conformations within the receptor extracellular domain (**Figure 4**). Close examination of these structures, and comparison to the GLP-1 bound structure, provides mechanistic insights into how PF 06882961, but not OWL-833 can closely mimic the pharmacology of GLP-1.

**Figure 4.**
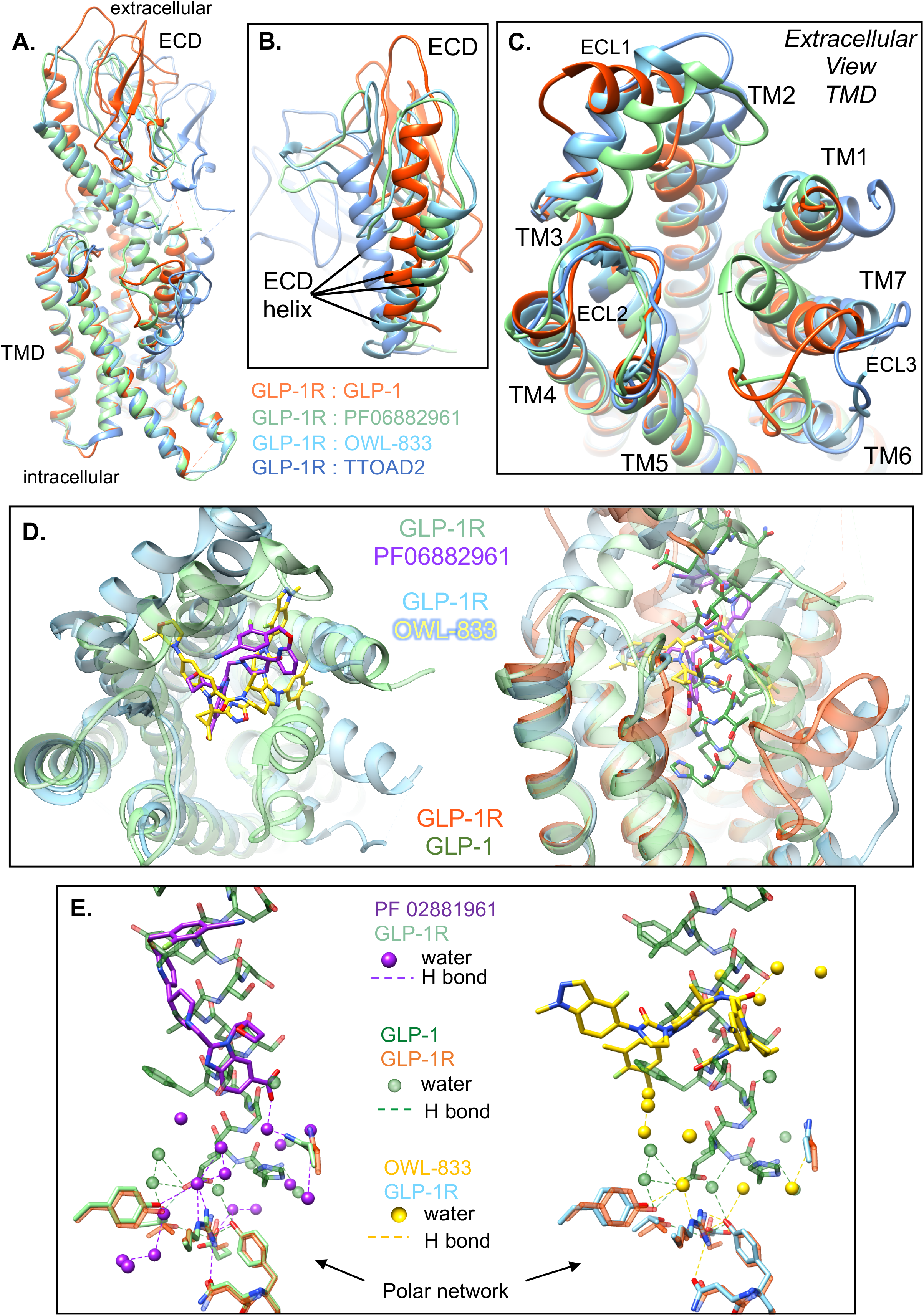
Comparisons of the GLP-1R conformations and binding pockets stabilised by GLP-1 and non-peptidic agonists. **A-C,** Superimposition of the receptor from the GLP-1R:G_s_ complex structures with GLP-1-(orange), PF 06882961- (pale green), OWL-833- (pale blue) and TT-OAD2-bound (6ORV – cornflower blue). **A,** Overlay of full length receptors; **B**; close up of the ECD showing the distinct location of the ECD alpha helix in the different structures; **C**, Close up of the GLP-1R TM bundle viewed from the extracellular side reveals differences in the conformations of TM1, TM2/ECL1, and the TM6-ECL3-TM7 backbones when stabilised by different agonists. **D,** Left, superimposition of the TMD from the PF 06882961- (pale green) and the OWL-833-bound (pale blue) receptors reveals partial, but limited overlap in the ligand binding site for PF 06882961 (purple) and OWL-833 (yellow); Right, side view of the overlaid upper portions of the TM bundles for GLP-1R bound to GLP-1 (receptor orange, GLP-1 - green), PF 06882961 (receptor pale green, ligand purple) and OWL-833 (receptor pale blue, ligand - yellow) reveals both non-peptide ligands overlap the binding of GLP-1, but stabilise different conformations in the TM bundle. **E,** PF 06882961 (purple) has a larger degree of overlap with GLP-1 (green) compared with OWL-833 (yellow). The extensive water network below the PF 06882961 binding site (purple spheres) overlap the N-terminus of GLP-1 and forms an extensive hydrogen bond network with polar residues deep in the TM binding cavity. The OWL-833 complex also contains structural waters (yellow spheres) that interact with polar residues deep in the GLP-1R TM cavity or with ligand. *See also Figures 5, 6, S5 and S6 and Video S1*.

OWL-833 exhibited a planar pose (parallel to the membrane) in the receptor TM bundle, with overlap with GLP-1 limited to residues F12^GLP-1^-L16^GLP-1^, with the OWL-833 fluoro methyl indazole and dimethyl morpholine arms extending beyond the pocket occupied by peptide and PF 06882961 (**Figure 4**). Despite this, the majority of side chains that form interactions with OWL-833, also interact with the peptide (**Figure 3, Table S2**) including P137^1.32^, E138^1.33^, L141^1.36^, L144^1.39^, Y145^1.40^, Y148^1.43^, K197^2.67^, L201^2.71^, Y205^2.75^, F230^3.33^, M233^3.36^, T298^ECL2^, N300^ECL2^ and L388^7.43^, albeit the nature of these interactions were very distinct and unlike GLP-1, OWL-833 did not form strong interactions with TM7. In contrast, PF 06882961 docked in a buried pocket within the receptor, adopting an elongated pose that overlapped the location of residues G10^GLP-1^-E21^GLP-1^ of GLP-1, and is principally located within the molecular envelope of GLP-1 in the peptide-bound structure (**Figure 4**). This leads to a subset of ligand interactions with the shared receptor residues, including S31^ECD^, L32^ECD^, L141^1.36^, K197^2.67^, L201^2.71^, Q221^ECL1^, F230^3.33^, M233^3.36^, T298^ECL2^, R380^ECL3/7.35^, L384^7.39^ (**Figures 3, S4 & Table S2**). Nonetheless, even within this region, GLP-1 has greater bulk that confers distinct interactions, including in engagement with ECL2, ECL1, TM7 and ECL3, which induce distinct receptor conformations (**Figure 4**). These structures raise the question; how does PF 06882961 closely mimic the pharmacology of the peptide, while OWL-833 only productively engenders a subset of these responses, specifically cAMP production?

Extensive GLP-1R alanine mutagenesis studies have revealed that ECL2 interactions and conformation are critical for all peptide-mediated G_s_-dependent cAMP signalling, while TM6/ECL3/TM7 are also important for this response, in a peptide-specific manner (Dods and Donnelly, 2015; Koole et al., 2012a; Koole et al., 2012b; Wootten et al., 2016c). In contrast, it is only the TM1/TM7/ECL3 domain that is critical for pERK1/2 that is a composite of *β*-arrestin-dependent and non-canonical G protein mediated signalling (Lei et al., 2018; Wootten et al., 2016c). Together with mutagenesis of residues deep within the receptor binding domain (**Figure S5**) (Graaf et al., 2016; Wootten et al., 2013b; Yang et al., 2016), this provides a framework for interpreting key features of the GLP-1R:G_s_ structures to understand characteristics linked to the common signalling of GLP-1 and PF 06882961, and to understand the distinct functional profile of OWL-833, and also of TT-OAD2 (through the recently reported GLP-1R structure (Zhao et al., 2020) with this compound).

Interestingly, while the overall conformation of ECL2, one of the most critical regions involved in G_s_-mediated signalling, is similar in the active GLP-1 and non-peptide-bound structures, there are distinctions in how this is maintained (**Figure 3**). In the GLP-1-bound structure, direct interaction between side chains that are located above and below ECL2 constrain the conformation of this domain. In contrast, there is limited direct interaction between PF 06882961 and ECL2, however, the conformation is instead constrained by water mediated interactions between ECL1, the ECD, particularly W33^ECD^, and ECL2, while water-mediated interactions within ECL2 also contribute to conformational stability. While the OWL-833 compound has a very distinct docking pose, ECL2 is stabilised by ligand interactions, as well as similar interactions between the ECD and ECL2 (**Figure 3**) that likely play a similar role in supporting G_s_-mediated signalling. In the previously reported TT-OAD2 structure, ECL2 also adopts a similar “active” conformation that would support G_s_-coupling and activation.

As noted above, potent canonical signalling by GLP-1 requires the far N-terminal dipeptide H7^GLP-1^-A8^GLP-1^ that resides deep within the receptor TM binding cavity and makes extensive interactions with receptor residues known to contribute to receptor activation. Remarkably, the PF 06882961 binding cavity extends below the ligand deep into the receptor core and there is overlap of this deep pocket with the binding site of the GLP-1 N-terminal H7^GLP-1^-E9^GLP-1^, and also the waters below the peptide that facilitate interaction with the central polar network of the receptor (**Figures 3 & 4E**). Moreover, the structural differences at the extracellular face of the receptor between the PF 06882961 and GLP-1 bound structures (discussed in detail below) ensure that the deep pocket is not exposed to bulk solvent in the PF 06882961-bound structure (**Video S1**). Consequently, there is a well-resolved structural water network that extends below the PF 06882961 binding site (**Figure 3**) that directly overlaps the location of H7^GLP-1^-E9^GLP-1^ and stabilises a similar conformational arrangement of the GLP-1R residues deep within the binding pocket, including those that interact directly, or via water, with GLP-1 (**Figure 4E**).

The recently published inactive structure of the full length GLP-1R (6LN2) (Wu et al., 2020), which likely closely correlates to the apo state, allows greater understanding of the transitions that occur within the GLP-1R as it moves from the inactive conformation to different active states (**Video S2**). In the inactive state, the GLP-1R ECD is stabilised in a “closed conformation” with the ECD peptide binding groove oriented towards the ECLs through interactions of the ECD N-terminal helix with ECL3 and the top of TM7, and the ECD loop region between S116-E127 with ECL1, with ECL1 itself providing an additional cap to the receptor core as well as forming interactions with ECL2 that likely constrain the conformation of this latter domain. The importance of the interaction between the far ECD and ECL3 for maintaining receptor quiescence is supported by mutagenesis and biochemical studies with GLP-1R and the related GCGR that reveal loss of this interaction can reduce the barrier to activation, or engender constitutive activity in the case of the GCGR (Wu et al., 2020; Yang et al., 2015; Yin et al., 2016; Zhao et al., 2016). Nonetheless, molecular dynamics simulations indicate that the ECD is dynamic and can transiently, partially or fully, disengage from the receptor core and this is likely key to both peptide and small molecule binding (Wu et al., 2020; Yang et al., 2015; Zhang et al., 2019; Zhao et al., 2020).

During receptor activation, there is release of the interactions between the ECD and receptor core and an upward and clockwise rotation of the ECD that is greatest upon accommodation of GLP-1 (**Figure 5A, Video S2**). In the small molecule-bound structures, this ECD translation is more limited and the far N-terminal helix caps the binding site for both PF 06882961 and OWL-833. In these conformations the ECD maintains interactions with the ECLs, particularly ECL1 and ECL2, in addition to the extreme ECD forming interactions with TM7 (PF 06882961) and TM1 (OWL-833) (**Figure 3**). There is also an outward movement of ECL1 that is enabled by the reorganisation of the TMD:ECD interactions for all ligands (**Figures 5**). For the non-peptidic agonists the extent of ECL1 translation is dictated by the ligand binding pose and the interactions of the ECD with TMs 1 or 7. In contrast to PF 06882961, the more planar pose of OWL-833 enables the dimethyl morpholine arm to extend towards ECL1 leading to a greater outward translocation of this domain relative to that seen in the PF 06882961-bound structure (**Figures 3 & 4D**). Nonetheless, in both small molecule-bound structures the conformation of ECL1 is similar to that observed in the inactive structure, whereas it undergoes reorganisation to accommodate peptide binding (**Figure 5, Video S2**). The greatest diversity across the GLP-1, PF 06882961 and OWL-833 bound structures is in the location and conformation of TM6/ECL3/TM7 and their interface with TM1 (**Figures 4 & 5**). While speculative, it is likely that disengagement of the interaction of the ECD N-terminal helix with ECL3 that is present in the “apo-like” structure, leads to increased mobility and outward movement of this domain, as reflected in inactive structures of class B1 GPCRs that lack the ECD, and in the TT-OAD2-bound structure where the ligand does not directly interact with this receptor segment. As such, ligands capable of forming direct interactions with the top of TM7/ECL3 lead to an overall contraction of the binding pocket relative to those that do not. The key structural differences in the positions and/or conformations of TM7, ECL1 and the ECD observed between the GLP-1 and PF 06882961, and indeed OWL-833 –bound, structures (**Figure 4 & 5**) contribute to both the ability of PF 06882961 to mimic the action of GLP-1, and the divergence in signalling and regulation seen with OWL-833.

**Figure 5.**
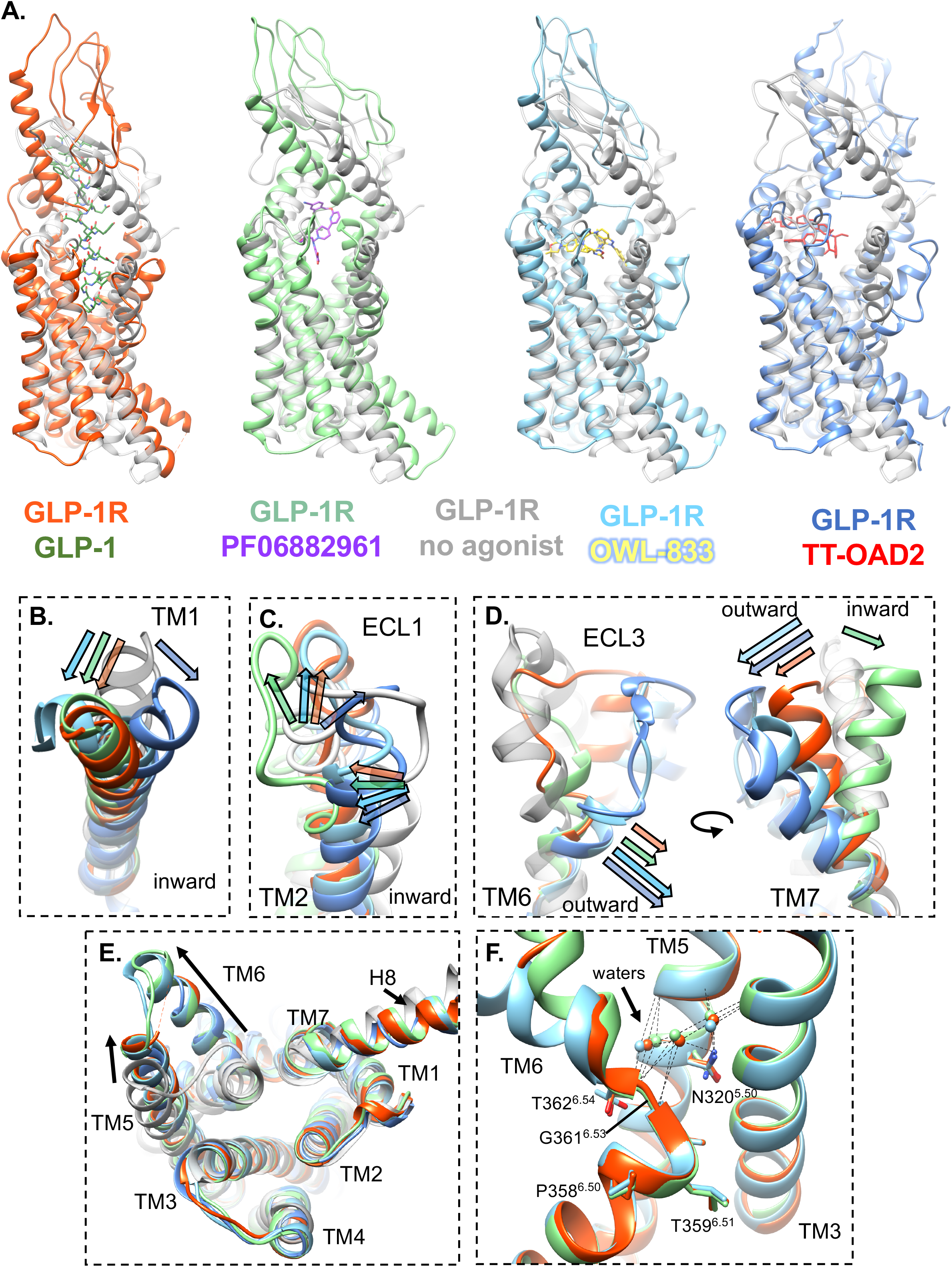
GLP-1R conformational transitions from the inactive conformation to different active states. **A,** Superimposition of the inactive ligand free GLP-1R structure (6LN2 – pale grey) with the GLP-1R bound to GLP-1 (orange), PF 06882961 (pale green), OWL-833 (pale blue) and TT-OAD2 (6ORV, cornflower blue) (from left to right). The ECD undergoes a large conformational transition upon binding of all agonists, but the degree of movement differs depending on the bound ligand. **B-D,** depicts the regions within the upper region of the TMD that undergo the largest movements when transitioning from the inactive receptor conformation to the different active states. TM1 and TM2 move inwards towards the centre of the TM bundle, and there is an upward movement of ECL1 for all agonists, however the exact nature of these movement differs (B-C). The largest differences are observed in TM6 and TM7 (D). The top of TM6 unwinds in the presence of all agonists, however only OWL-833 and TT-OAD2 promote a significant outward movement. Large outward movements within ECL3 and TM7 are also observed upon binding of OWL-833 and TT-OAD2, and to a lesser extent GLP-1, however PF 06882961 promotes a significant inward movement (D). **E**, At the intracellular face, all agonists promote the same receptor conformation that involves a large outward movement of TM6 and a smaller outward movement of TM5. There is also a small downward movement of H8 away from the detergent micelle. **F,** The TM6 kink around the conserved P^6.50-^X-X-G^6.53^ motif is stabilised by a similar network of residues in all active state structures and is also stabilised by structural waters (spheres depict waters coloured relative to their bound ligand (orange GLP-1, pale green PF 06882961, pale blue OWL-833), with hydrogen bonds depicted by dotted lines. *See also Figures 4, S5 and S6 and Video S2*.

In the PF 06882961 bound structure, TM7 is translated dramatically inwards, overlapping with the molecular envelope of GLP-1. This both stabilises compound binding and seals the deeper pocket from access to bulk water (**Figures 3 & 4, Video S1**). Intriguingly, despite the large contraction of the upper part of TM7 induced by the binding of PF 06882961, this region along with the structural waters enables interaction networks within the receptor that can mimic the active receptor conformation induced by GLP-1 (**Figures 3, 6**). In the GLP-1-bound structure, the more outward conformation of ECL3 and TM7 is stabilised by an extensive polar network that extends from D15^GLP-1^ of the peptide that forms a salt bridge with R380^ECL3/7.35^. From this ligand interaction, receptor residues R380^ECL3/7.35^, D372^ECL3^, W306^5.36^, R310^5.40^ and E373^ECL3^ form consecutive interactions via a salt bridge, hydrogen bond, cation-π stacking and salt bridge, respectively, where all but E373^ECL3^ also form interaction with the GLP-1 peptide (**Figure 6A**). This latter residue however, is important for the differential engagement of GLP-1 and the biased agonist peptide, ExP5, demonstrating importance of the ECL3 conformation and signalling efficacy (Zhao et al., 2020). Furthermore, the ECL3 interaction network interconnects with the water network that stabilises the N-terminus of GLP-1 (H7^GLP-1^-E9^GLP-1^) and the central polar core of the receptor, which is mediated via K383^7.38^ (**Figures 3 & 6A**). Alanine mutation of R310^5.40^, K383^7.38^, D372^ECL3^ and E373^ECL3^ reduce the potency of GLP-1 by over 1,000-fold in multiple signalling pathways (cAMP, calcium and pERK1/2), while these mutations have a much smaller effect on affinity (Coopman et al., 2010; Wootten et al., 2016b; Wootten et al., 2016c). Together, the structural and mutagenesis studies support the strong influence of this polar interaction network in GLP-1-mediated signal transduction and stabilisation of the overall active conformation.

**Figure 6.**
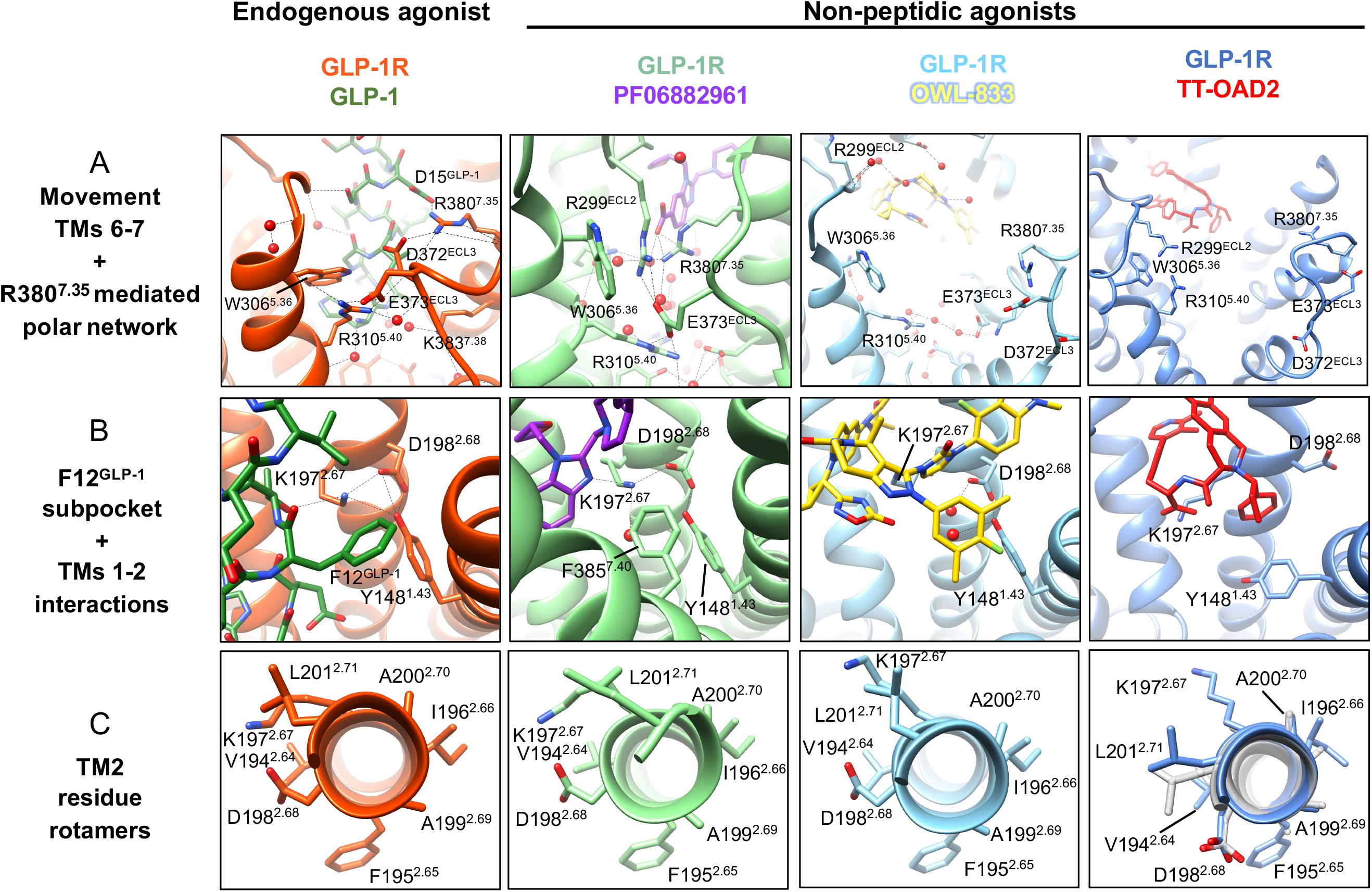
Distinct interaction networks provide a structural rationale for how PF 06882961, but not OWL-833, can mimic the signalling of GLP-1. **Top,** The conformation of TM6/ECL3/TM7 in the GLP-1-(far left) and PF 06882961-bound (middle left), but not the OWL-833 and TT-OAD2 bound (right panels) GLP-1Rs, enable the formation of an extensive polar interaction network. This is mediated by interactions of R380^7.35^ with D15^GLP-1^ of the peptide or the carboxylic acid of PF 06882961. In the PF-06882961, this network stabilises the water mediated network below the bound ligand that overlaps the peptide binding site. **Middle,** The TM7 contraction induced by PF 06882961 moves F385^7.40^ into a hydrophobic groove between TM1 and TM7 (middle left), occupying the same pocket filled by F12^GLP-1^ in the peptide bound structure (far left), thereby stabilising common overall rearrangements within side chains of key residues located in TMs 1 and 2. The fluoro dimethyl phenyl moiety of OWL-833 plays a similar role in the OWL-833 bound structure (middle right), albeit different interactions of the ligand with K197^2.67^ stabilise a different conformation of this side chain. TT-OAD2 cannot fill this pocket and does not form this polar network (far right). **Bottom,** In the absence of the polar network, residues in TM2 adopt a different rotameric arrangement in the TT-OAD2 bound receptor, which is close to that observed in the inactive receptor - grey) when compared with GLP-1, PF 06882961 and OWL-833. *See also Figures 4, S3, S5 and S6*.

Together with interactions directly with the bound PF 06882961 scaffold, the inwardly translated TM7 conformation is able to mimic the polar interaction network observed in the GLP-1-bound structure (**Figures 3 & 6A**). Here, the carboxylate of PF 06882961 mimics D15^GLP-1^ of GLP-1 as they both share an ionic interaction with R380^ECL3/7.35^. The inward movement of TM7 is followed by TM6 enabling E373^ECL3^ to form the same salt bridge with R310^5.40^ that was formed in the GLP-1 complex (**Figure 6A**). These changes allow other shared polar residues to move and partake in the very extensive water-supported polar network that links the bound ligand to the central polar network (**Figure 3 & 4**). This includes, R299^ECL2^ that moves inward to form a salt bridge with E373^ECL3^ and cation-π stacking with W306^5.36^ thereby aiding the stabilisation of TM6 and TM7 in the contracted conformation of the vestibule (**Figure 6A**). In support of this, alanine mutation of R380^ECL3/7.35^ or E373^ECL3^ had a large impact on cAMP production mediated by both GLP-1 and PF 06882961 (**Figure S5**). Furthermore, the TM7 contraction induced by PF 06882961 moves F385^7.40^ into a hydrophobic groove between TM1 and TM7 to occupy the pocket filled by F12^GLP-1^ in the peptide bound structure, thereby stabilising common overall rearrangements of side chains of key residues located in TMs 1 and 2 (**Figures 3 and 6B**). Both F385^7.40^ in the PF 06882961 structure and F12^GLP-1^ of GLP-1 show T-shaped π-stacking interactions with Y148^1.43^, which undergoes a rotameric change relative to inactive structures, leading to a hydrogen bond with D198^2.68^. D198^2.68^, in turn, forms a salt bridge with K197^2.67^ and these three residues are important for ligand interactions and activation as shown by large potency changes in mutational studies (**Figure S5**) (Coopman et al., 2010; Lei et al., 2018; Lopez de Maturana and Donnelly, 2002). These interactions also influence the rotamers of surrounding residues (**Figure 6C**), which either engage with peptide or with the water-network below the PF-06882961 binding site (**Figure 3**). The importance of F385^7.40^ in PF 06882961 activity was confirmed by alanine mutagenesis where this mutation reduced PF 06882961 potency by 10-fold with no influence on GLP-1 cAMP production (**Figure S5**). The similarities in the interaction networks formed within the GLP-1- and PF 06882961-bound receptor result in similarities in the electrostatic and hydrophobic properties of their binding pockets, despite differences in their overall shape and volume (**Figure S6, Video S1**).

OWL-833 is a potent agonist of the GLP-1R for cAMP production, albeit with lower efficacy than PF 06882961, and this contrasts with TT-OAD2 that is a low efficacy agonist for this pathway (Zhao et al., 2020) (**Figure 7**). The fluoro methyl indazole arm of OWL-833 extends towards TM1 and TM2, with the fluoro dimethyl phenyl moiety π-stacking with Y148^1.43^ in a similar manner to F12^GLP-1^, and F385^7.40^ in the PF 06882961 structure, and supporting the polar interactions with and within TM2. By comparison, while there is considerable overlap with the pose of OWL-833, TT-OAD2 lacks an aromatic extension equivalent to the fluoro dimethyl phenyl group of OWL-833, and Y148^1.43^ and D198^2.68^ adopt distinct rotameric positions compared to the other structures and cannot form an equivalent polar network to constrain the TM1-TM2 interface (**Figure 6B**). As these residues play a critical role in ligand potency of the other small molecule agonists, it is likely that the observed conformational differences contribute to the lower efficacy of TT-OAD2.

**Figure 7.**
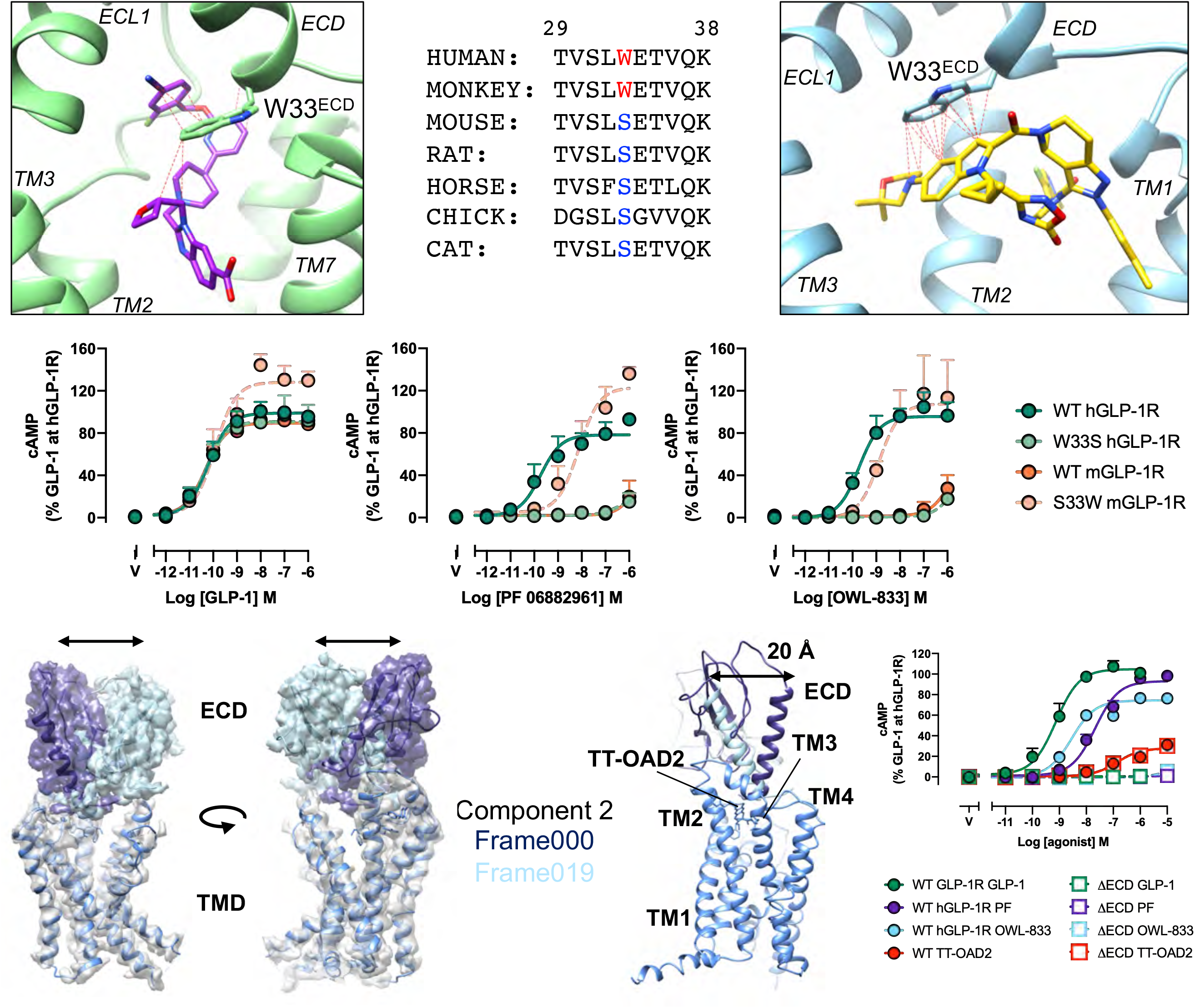
The species selectivity of PF 06882961 and OWL-833, and the importance of the ECD in GLP-1, PF-06882961, OWL-833 and TT-OAD2 activity can be rationalised by the cryo-EM structures. **Top,** PF 06882961 (top left) and OWL-833 (top right) form extensive contacts with W33^ECD^ within the ECD of the GLP-1R. This residue is only conserved in the human and primate GLP-1R, but not in rodents and other species (top middle). **Middle,** concentration response curves for cAMP production for GLP-1 (middle left), PF 06882961 (middle) and OWL-833 (middle right) at the human (h)GLP-1R and mouse (m)GLP-1R reveal that PF 06882961 and OWL-833 are not active at the mouse GLP-1R. Swapping the residue at position 33 between hGLP-1R and mGLP-1R (W33S hGLP-1R and S33W mGLP-1R) switches the species selectivity. **Bottom,** 3D variability analysis using cryoSPARC: left orthogonal views of the cryo-EM maps from Frames 000 and 019 from component 1 show extensive mobility in the ECD relative to the TM bundle of the TT-OAD2 bound GLP-1R. Measurement of the distance between the Cα of E52 (of ECD models rigid body fit to the two extreme maps) revealed a 20 Å movement of the ECD. Bottom right, cAMP production concentration-response curves for GLP-1 (green), PF 06882961 (purple), OWL-833 (pale blue) and TT-OAD2 (red) at the full length GLP-1R (closed circles) and the “headless” GLP-1R that lacks the ECD (open squares). This shows the ECD is not required for the partial agonism exhibited by TT-OAD2, but is essential for the activity of GLP-1, PF 06882961 and OWL-833. Pharmacological data are presented as the mean + S.E.M of at least 3 independent experiments performed in triplicate or duplicate. *See also Figure S5 and Videos S3 and S4*.

In contrast to the GLP-1 and PF 06882961-bound GLP-1R structures, there is only very limited interaction of OWL-833 with TM7. This leads to an outwardly splayed conformation of TM7/ECL3/TM6 that is similar that observed in the TT-OAD2-bound structure (**Figure 4A, C**). This position of TM6/ECL3/TM7 exposes the deep binding pocket below OWL-833 to bulk water (**Figures 3 & 6A**). Density in the cryo-EM map corresponding to waters was present deep in the OWL-833 binding pocket, where they stabilise the rotameric positions of the deep central polar network of the receptor (**Figure 4E**). However, unlike the PF 06882961-bound structure, these did not form a continuous network extending to the ligand (**Figures 3 & 4E**), albeit additional weak density that could not be confidently assigned to waters was also present in the cryo-EM map. Despite the overall higher resolution within the ligand binding site, the less well resolved water density of the OWL-833 binding pocket compared to the PF 06882961 bound map is consistent with greater mobility of waters in the larger binding cavity, and is likely a result of the lack of the extensive polar interactions extending from TM7 that were observed in the PF 06882961 bound complex (**Figure 6A**). These differences also lead to distinctions in the hydrophobicity and electrostatic potential of the deeper binding cavity of the OWL-833 complex (**Figure S6A**). These differences in the hydrogen bond network within TM7-TM6-TM5 may account for the lower efficacy for the cAMP pathway observed with OWL-833, relative to PF 06882961. Indeed, alanine mutation of E373^ECL3^, one of the key mediators of this network in the GLP-1 and PF-06882961 bound structures, had a much more limited influence on OWL-833-, when compared to PF 06882961- and GLP-1-mediated cAMP production (**Figure S5**).

Mutagenesis studies have demonstrated that TM1 and TM6-ECL3-TM7 plays an important role in biased agonism of peptide agonists (Lei et al., 2018; Wootten et al., 2016c). This is further supported by the structures of the GLP-1R bound to OWL-833, TT-OAD2 and the biased peptide, ExP5 (Liang et al., 2018b; Zhao et al., 2020). In this latter structure, the TM6-ECL3-TM7 region resides in an intermediate position to that induced by GLP-1 and by OWL-833 and TT-OAD2 (**Figure S6C**). ExP5 is a G protein-biased peptide relative to GLP-1 with greater coupling to cAMP and calcium mobilisation relative to *β*-arrestins and receptor internalisation, where it is only a partial agonist (Zhang et al., 2015). Thus, the relative positioning of the TM6-ECL3-TM7 domain to the rest of the bundle is correlated to the ability to couple to regulatory pathways with a more open conformation resulting in reduced responses, and providing a structural basis for the differences in the signalling and regulatory profile of OWL-833 (and TT-OAD2 and ExP5) relative to GLP-1 and PF 06882961.

Despite differences in the extracellular conformation induced by GLP-1 and non-peptide agonists, these all converge to similar conformational transitions at the base of the GLP-1R binding pocket, where all non-peptide agonists and GLP-1 promote a similar reorganisation of polar residues that enables the formation and stabilisation of the TM6 kink, which is a hallmark of the active state of class B GPCRs, and includes structural waters (**Figure 5**). While the resolution of the published TT-OAD2 bound structure did not allow for the visualisation of waters (Zhao et al., 2020), the almost identical rotameric positioning of these side chains suggest the same stabilisation networks exist. At the intracellular face of the receptor, all agonist-bound structures adopt an equivalent conformation with the most predominant feature being a large outward movement of TM6 relative to the inactive receptor, consistent with previous observations in active state structures of class B1 GPCRs (**Figure 5**) (Liang et al., 2020a; Liang et al., 2020b; Liang et al., 2018a; Liang et al., 2018b; Liang et al., 2017; Ma et al., 2020; Qiao et al., 2020; Zhang et al., 2017; Zhao et al., 2019; Zhao et al., 2020). As all the structures are coupled to G protein, it is likely that the common interaction networks in the centre and base of the receptor, regardless of the agonist, are allosterically influenced by the presence of the bound G protein. The interactions formed between the G_s_ protein and the receptor were equivalent between the GLP-1, PF 06882961 and OWL-833 bound complexes and consistent with previously published structures of agonist bound GLP-1R:G_s_ complexes (Liang et al., 2018b; Zhang et al., 2017; Zhao et al., 2020), albeit that additional interactions (including water-mediated interactions) were identified in the current study due to the higher resolution of the data (**Figure S7**).

Collectively, these data reveal structural similarities and differences associated with GLP-1 and PF 06882961 binding that provide mechanistic insight into how PF 06882961 mimics key conformational features of the GLP-1-bound receptor, which enable similar signalling profiles by these very distinct classes of agonists, while providing confirmatory evidence for the role of the TM1/TM7/ECL3/TM6 domain in engendering biased agonism.

### Receptor and G protein dynamics contribute to agonist-dependent G protein coupling and activation

The 2D projections from the cryo-EM imaging include particles that span 3D conformational space present at vitrification and therefore provide an avenue to resolve dynamics from static image data. 3D variability analysis in cryoSPARC (Punjani et al., 2017) was used to extract the 3 major principal components that contributed to conformational variance for each of the structures. This revealed common, coordinated twisting and rocking motions of the receptor, relative to the G protein, for all 3 structures (**Video S3**). Notably, equivalent analysis of the previously solved TT-OAD2 structural data suggested that this complex has a more stable receptor-G protein interface with reduced relative motion compared to the other structures (**Video S3**). Moreover, in contrast to the complexes of the more efficacious agonists for G_s_ activation, the *α*-helical domain (AHD) of G*α*_s_ in the TT-OAD2 bound structure only occupies a single, open, orientation, albeit that this was poorly resolved, indicative of substantive motion within this location (**Video S4**). For each of the other complexes, the AHD undergoes translational motion between open and more closed orientations, relative to the G*α*_s_ Ras-like domain (**Video S4**). Collectively, the predominant open conformation of G*α*_s_, and more limited dynamics of the receptor-G protein interface in the TT-OAD2 structure are consistent with the slow G_s_ activation kinetics and rate of cAMP production previously observed for this ligand, though additional experiments are required to better understand the unique pharmacological behaviour of TT-OAD2 compound.

Due to the conformational variance observed in the location of the G*α*_s_ AHD in the GLP-1, PF 06882961 and OWL-833 bound complexes, we performed additional G protein focused refinements that enabled the identification of five distinct classes with differences in the AHD position for each complex (**Figure S7**). Three classes were common to all three structures (labelled up (most open), middle, and down (most closed)), although the number of particles present in each class differed for the three agonists, suggestive of different mobilities and conformational ensembles stabilised by each ligand. A morph between these conformations is shown in **Video S4**. While not the most populated class, the “down” AHD position was the best resolved and the only class that allowed confident modelling into the map (**Figure S7**). The other 2 classes were more varied and appeared to only partially overlap across the different structures. While the distinct sampling of AHD position across the structures would also be consistent with each of the agonists having discrete efficacy, it is important to note that the biochemical system used for complex formation is artificial (detergent micelle, dominant negative G*α*_s_, Nb35) and that extrapolation of observed behaviour to the normal cellular membrane environment needs to done with a high-degree of caution.

In-line with the lower resolution of ECL1 and the receptor ECD in consensus maps of the GLP-1 bound receptor, these domains exhibited greater motion, relative to the rest of the bundle, compared to the equivalent domains in the PF 06882961 and OWL-833 bound receptors (**Figure S2, Videos S3 & S4).** Modelling the receptor backbone into the two extremes revealed a coordinated 7 Å movement in these regions relative to the remainder of the bundle, when overlaid in the maps from the two extreme frames of the analysis (**Video S2**). This also revealed a 4.5 Å movement of the location of the C-terminus of the GLP-1 peptide, while the N-terminal region that binds deep within the TM bundle and is responsible for activation, is relatively stable. Interestingly, while ECL1 is relatively stable, the ECD of the TT-OAD2 bound structure is extremely mobile with more substantial motions than those induced by GLP-1, transitioning 20 Å between the two extremes when measured from the C*α* of E52^ECD^ in models built into these maps (**Figure 7, Video S3**). This mobility is also reflected in the low resolution of the GLP-1R ECD in the consensus map of the complex with TT-OAD2 (Zhao et al., 2020). This substantial flexibility is likely related to the lack of stabilising interactions formed between the ligand with the ECD, and a lack of interaction of the ECD with ECLs1/2 and TMs1/7 that were present in the PF-06882961 and OWL-833 structures. The ECD of the GLP-1R has been identified in previous studies to contribute to agonist-induced cAMP signalling (Yin et al., 2016). The ability of PF 06882961 and OWL-833 to engender a stable ECD conformation more akin to that of peptide agonists, when compared to TT-OAD2, may also contribute to their higher efficacy in cAMP production. Interestingly, the ECD is crucial for the function of GLP-1, PF 06882961 and OWL-833, but is not required for the cAMP signalling mediated by TT-OAD2 as revealed from cAMP studies using a GLP-1R construct lacking the N-terminal ECD (**Figure 7**).

### PF 06882961 and OWL-833 have species selectivity that can be explained by their unique interaction patterns relative to peptide agonists

PF 06882961 and OWL-833 form substantial hydrophobic interactions with W33^ECD^ within the N-terminus of the receptor ECD that are not observed with GLP-1 (**Figure 7, Table S2**). Mutation of W33 to alanine completely abolished PF 06882961 and OWL-833-induced cAMP production, with no effect on GLP-1-mediated signalling (**Figure S5**). Tryptophan at position 33 is only conserved in human and primate GLP-1R, but not in other mammalian species where residue 33 is a serine. In cell lines expressing rodent receptors, OWL-833 and PF 06882961 were unable to promote cAMP production revealing that these compounds display species selectivity (**Figure 7**). Mutation of W33^ECD^ to serine in human GLP-1R had a similar effect to that observed with alanine mutation, abolishing cAMP signalling, however a serine to tryptophan mutation in rodent receptors was sufficient to recover activity of these compounds (**Figure 7**), confirming the essential role of W33^ECD^ in determining this species selectivity. This has significant implications for preclinical development of GLP-1R small molecule agonists, as these are often assessed in rodent models of type 2 diabetes and obesity.

## CONCLUDING REMARKS

Diabetes and obesity are major health burdens and the GLP-1R has been validated as a target to regulate metabolism for treatment of these diseases(Brown et al., 2018; Nauck and Meier, 2019). While there is major interest in the development of non-peptide drugs for the GLP-1R that may be orally delivered, this has proven profoundly difficult and only three compound series have progressed into clinical development. We have identified unique activation profiles for each of these non-peptide ligands and differences in how these compounds bind to and activate the GLP-1R. The cryo-EM structures provide a deeper understanding of the active state conformations that the GLP-1R is capable of adopting and reveal new druggable pockets for the development of small molecule agonists. The near-atomic level detail revealed within the binding pockets induced by GLP-1, PF 06882961 and OWL-833 provide critical information that can be utilised in structure-guided-activity programs to progress development of the next generation of non-peptide agonists for this receptor. Moreover, PF 06882961 is able to better mimic the in vitro pharmacological actions of GLP-1 than some closely related peptides, making this a particularly attractive lead structure, and this activity can be rationalised from the new high-resolution structures. In particular, the visualisation of the water-mediated interaction network below the ligand binding site, and how this is stabilised by the unique interaction patterns with PF-06882961, provides a novel template for the design of new agonists that could extend out and access this water pocket. This could potentially provide higher affinity/efficacy compounds than those currently available. Furthermore, the spectrum of non-peptide agonist binding sites identified herein for the GLP-1R may extend to other class B1 GPCRs that are of high therapeutic potential. These receptors are likely capable of adopting similar conformations, if targeted by appropriate small molecule scaffolds, albeit the specific nature and properties within these pockets may differ between receptors due to their differing sequences. Our work opens up new avenues to pursue small molecule agonists identification and development.

## Acknowledgements

This work was supported by the National Health and Medical Research Council of Australia (NHMRC) (project grant 1126857, ideas grant 1184726 and SRF 1160076 (D.W.)), and program grant 1150083 (P.M.S.)). P.M.S. is a Senior Principal Research Fellow and D.W. a Senior Research Fellow of the NHMRC. R.D. was supported by Takeda Science Foundation 2019 Medical Research Grant and Japan Science and Technology Agency PRESTO (18069571). D.E.G. was funded by the Lundbeck Foundation (R163-2013-16327). A.J.K acknowledges financial support from the Independent Research Fund Denmark (8021-00173B). This work was partially supported the Monash University Ramaciotti Centre for cryo-electron microscopy, and the Monash University MASSIVE high-performance computing facility.

## Author Contributions

**Conceptualization**: D.W. and P.M.S. conceived the project and formulated the research plan. **Investigation**: X.Z. expressed and purified receptor complexes; R.D., M.B., H.V. performed sample vitrification and cryo-EM imaging; X.Z., M.B., R.D. processed the EM data; X.Z., M.B. generated atomic models; X.Z., P.Z., T.T.T., S.Y.A., G.D.S., C.R.U., T.E. generated DNA constructs and performed pharmacological assays. **Formal Analysis**: D.W., P.M.S., A.J.K., D.E.G., X.Z., M.B. analysed the receptor structures; D.W., X.Z, P.Z, T.T.T, S.Y.A., G.D.S., C.R.U., S.R-R. analysed pharmacological data. **Resources**: S.G.B.F generated FLAG affinity resin; P.S synthesised OWL-833. **Supervision**: D.W. and P.M.S supervised the whole project; A.G and Y-L.L assisted with supervision for complex biochemistry and molecular modelling; P.Z, S.R-R, C.J.L supervised pharmacological experiments. **Project administration**: D.W. and P.M.S coordinated the biochemistry and pharmacological studies, data processing and analysis; R.D coordinated sample vitrification and cryo-EM imaging; S.R-R coordinated the species selectivity analysis. **Visualization**: D.W., X.Z., M.B., A.J.K., R.D., P.M.S., prepared figures and videos for the manuscript. **Writing - original draft:** D.W., P.M.S., X.Z., A.J.K and D.E.G wrote/contributed to the original manuscript draft. **Writing - review & editing:** M.B, P.Z, T.T.T, S.Y.A, C.R.U, A.C, S.G.B.F, L.J.M, S.R-R, C.J.L, R.D contributed to writing, review and editing of the manuscript. **Funding Acquisition**: D.W., P.M.S., R.D., D.E.G acquired the financial support for the project leading to this publication.

## Declarations of interest

C.R.U, T.E and S.R-R are employees of Novo Nordisk. P.S is an employee of Apigenex.

## STAR METHODS

### CONTACT FOR REAGENT AND RESOURCE SHARING

Further information and requests for resources and reagents should be directed to and will be fulfilled by the Lead Contact, Denise Wootten.

### EXPERIMENTAL MODELS

Protein expression for biochemical analysis and purification was performed in *Spodoptera frugiperda (Sf9)* and *Trichuplusia ni* (*Tni*) insect cells, maintained in ESF 921 serum free media (Expression systems) at 30°C. CHOFlpIn cells used for mammalian cell studies were maintained in DMEM supplemented with 5% FBS at 37°C in 5% CO_2._ *Escherichia Coli* (*E. coli*) strain BL21, used to express Nb35, was cultured in Luria-Bertani (LB) liquid medium (10 g tryptone, 10 g NaCL and 5 g yeast extract per litre) with continuous shaking (180 rpm) or on LB agar plate (LB medium with 15 g agar per litre) at 37°C.

## METHOD DETAILS

### Constructs

The human GLP-1R was modified to replace the native signal peptide with that of haemagglutinin (HA) to improve receptor expression and contain an N-terminal Flag tag and C-terminal 8xHis tag. 3C protease cleavage site (LEVLFQGP) was inserted between both tags and the receptor. The constructs were generated in both mammalian (pcDNA3.1) and insect cell expression (pFastBac) vectors, and the modification did not affect the receptor pharmacological profile (Liang et al., 2018b). Dominant negative Gαs (DNGαs) was used has been previously described (Liang et al., 2020b; Liang et al., 2018c). 8xHis tagged Nb35 was provided by B. Kobilka (Rasmussen et al., 2011). Various constructs were used for mammalian cell assays including human GLP-1R in pcDNA3.1 modified with a 2xcMyc-N-terminal epitope tag, and human GLP-1R with a c-terminal Rluc-8 tag. These constructs were validated previously (Fletcher et al., 2018; Koole et al., 2010). For G protein conformational assays, a Nanoluc tag was inserted into Gαs after G(h1ha10) with a SGGGGS linker as described previously (Liang et al., 2018c). This was used in conjunction with Gβ1 and an N-terminally Venus labelled Gγ2.

### Insect cell expression

The Bac-to-Bac Baculovirus Expression System (Invitrogen) was used to generate high-titre recombinant baculovirus (> 10^9^ viral particles per ml). Recombinant baculovirus was produced by transfecting recombinant bacmids (2-3 μg) into *Sf9* cells (5 mL, density of 4×10^5^ cells per mL) using FuGENE HD Transfection Reagent (Promega) and Opti-MEM Reduced Serum Media (Thermo Fisher Scientific). After 5 d incubation at 27°C, P0 viral stock was harvested as the supernatant of the cell suspension to produce high-titre viral stock (P1 and P2 virus). Viral titres were analysed by flow cytometry on cells stained with gp64-PE antibody (Expression Systems). Human GLP1R, human DNGαs and human Gβ1-γ2 were co-expressed by infecting *Tni* insect cells at a density of 3.5 million/mL with P2 baculovirus at multiplicity of infection (MOI) ratio of 3:3:1. Culture was harvested by centrifugation 48 h post infection and cell pellets were stored at −80°C.

### Purification of Nanobody35

Nanobody-35 (Nb35) bearing a C-terminal His-tag was expressed in the periplasm of *E. coli* strain BL21 following induction with IPTG. Cultures of 1L were grown to OD_600_ to 0.7 at 37 °C in TB media containing 0.1% glucose, 2 mM MgCl_2_ and 100 μg/mL ampicillin. Induced cultures were grown overnight at 25 °C. Cells were harvested by centrifugation and lysed in ice-cold buffer (50 mM Tris pH 8.0, 0.125 mM sucrose, 0.125 mM EDTA). Lysate was centrifuged to remove cell debris and Nb35 was purified by nickel affinity chromatography (Pardon et al., 2014). Elute was concentrated to 5 mg/mL and stored at −80°C.

### Complex formation and purification

GLP-1R-DNGs complex formation and purification were performed as described by Liang, *et al*. (Liang et al., 2018b). Cell pellets (from 1 L insect cell culture) were thawed and suspended in 20 mM HEPES pH 7.4, 50 mM NaCl, 5 mM CaCl_2_, 2 mM MgCl_2_ supplemented with cOmplete Protease Inhibitor Cocktail tablets (Sigma Aldrich) and benzonase (Merk Millipore). Complex was formed by adding GLP-1R agonists (10 µM GLP-1, 50 µM PF-06882961 or 50 µM OWL833), Nb35–His (10 µg/ml) and apyrase (25 mU/ml, NEB). The suspension was incubated for 1 h at room temperature. The complex was solubilized from membrane using 0.5% (w/v) LMNG and 0.03% (w/v) CHS (Anatrace) for 1 h at 4°C. Insoluble material was removed by centrifugation at 30,000 g for 20 min and the solubilized complex was immobilized by batch binding to M1 anti-Flag affinity resin in the presence of 5 mM CaCl_2_. The resin was packed into a glass column and washed with 20 column volumes of 20 mM HEPES pH 7.4, 100 mM NaCl, 2 mM MgCl_2_, 5 mM CaCl_2_, 1 µM agonist, 0.01% (w/v) LMNG and 0.006% (w/v) CHS followed by elution with buffer containing 5 mM EGTA and 0.1 mg/ml FLAG peptide. The complex was then concentrated using an Amicon Ultra Centrifugal Filter (MWCO, 100 kDa) and subjected to size-exclusion chromatography (SEC) on a Superdex 200 Increase 10/300 column (GE Healthcare) that was pre-equilibrated with 20 mM HEPES pH 7.4, 100 mM NaCl, 2 mM MgCl_2_, 1 µM agonist, 0.01% (w/v) MNG and 0.006% (w/v) CHS to separate complex from contaminants. Eluted fractions consisting of receptor and G protein complex were pooled and concentrated to 3-5 mg/mL. The complex samples were flash frozen in liquid nitrogen and stored at −80°C.

### SDS-PAGE and Western blot analysis

Samples from important steps during purification were collected and analysed by SDS–PAGE and western blot. TGX™ Precast Gel (BioRad) was used to separate proteins within samples at 200 V for 30 min. Then gels were either stained by Instant Blue (Sigma Aldrich) or immediately transferred to PVDF membrane (BioRad) at 100 V for 1 h. The proteins on the PVDF membrane were probed with two primary antibodies simultaneously, rabbit anti-Gs C-18 antibody (cat. no. sc-383, Santa Cruz) against Gα_s_ subunit and mouse poly-His antibody (cat. no. 34660, QIAGEN) against His tags. The membrane was washed and incubated with secondary anti-mouse and anti-rabbit antibodies (LI-COR). The membranes were imaged on a Typhoon 5 imaging system (Amersham).

### Negative staining and data processing

The complex samples were diluted to 0.006 mg/mL in 20 mM HEPES pH 7.4, 100 mM NaCl, 2 mM MgCl_2_ and 1 µM agonist and applied to the continuous carbon grids (EMS). The grids were stained with 0.8% (w/v) uranyl formate solution and imaged using a cryo-EM Tecnai™ T12 or L120c TEM at 120 kV. Around 50 images for each complex were collected with a magnified pixel size of 2.06 Å, and ∼20,000 particles of each complex were auto picked, extracted and 2D classified by RELION-3.1-beta.

### Vitrified sample preparation and data collection

Samples (3 µL) were applied to a glow-discharged UltrAuFoil R1.2/1.3 Au 300 gold foil grids (Quantifoil GmbH, Großlöbichau, Germany) and were flash frozen in liquid ethane using a Vitrobot mark IV (Thermo Fisher Scientific, Waltham, Massachusetts, USA) set at 100% humidity and 4°C for the prep chamber and 10 s blot time. Data were collected on a Titan Krios G3i microscope (Thermo Fisher Scientific) operated at an accelerating voltage of 300 kV with a 50 μm C2 aperture at an indicated magnification of 105K in nanoprobe EFTEM mode. Gatan K3 direct electron detector positioned post a Gatan BioQuantum energy filter (Gatan, Pleasanton, California, USA), operated in a zero-energy-loss mode with a slit width of 25 eV was used to acquire dose fractionated images of the samples with a 100 µm objective aperture. Movies were recorded in hardware-binned mode (previously called counted mode on the K2 camera) with the experimental parameters listed in Table S1 using a 9-position beam-image shift acquisition pattern by custom scripts in SerialEM (Schorb et al., 2019).

### Cryo-EM data processing

5739 movies from the GLP1 dataset were motion corrected using MotionCor2 (Zheng et al., 2017) and subjected to contrast transfer function (CTF) estimation using Kai Zhang’s GCTF software (Zhang, 2016). 2.7 million particles were reference-free picked in RELION 3.0 (Zivanov et al., 2018), extracted with a 128 pixels box and subjected to 2D classification. An initial 3D model was generated from selected good 2D classes by Stochastic Gradient Descend (SGD) in RELION and was used for 3D refinement of the particle set. The resulting 3D map, low-pass filtered to 10 Å, was used to reference-based re-pick the micrographs in RELION, producing 3.8 million particles. The re-picked particles were extracted with a 128 pixels box (1.66 Å/pixel) and were subjected to two rounds of 3D classification with 3 classes, each time selecting the best-resolved class. The resulting set of 715K particles was subjected to three rounds of CTF refinement, Bayesian polishing with a larger box/smaller pixel and 3D auto refinement, followed by another round of 3D classification and 3D refinement that produced a 3D map at 2.14 Å in a 320 pixels box (0.83 Å/pixel) from a set of 636K particles. The particle set was then subjected to CTF refinement with higher order aberrations in RELION 3.1 that resulted in the final consensus map at 2.11 Å. The receptor and ECD were further refined with local masks in RELION.

5625 and 5859 movies of PF-06882961- and OWL833-GLP-1R-Gs complexes, respectively, were motion corrected using MotionCor2 (Zheng et al., 2017) and subjected to CTF estimation with GCTF software (Zhang, 2016). Particles were picked from corrected micrographs using crYOLO software (Wagner et al., 2019). Picked particles (2.36 million and 5.32 million particles of PF-06882961 and OWL833 complex respectively) were extracted using a box size of 288 pixels and 2D classified using RELION (version 3.1). The selected 2D particles (1.34 million and 2.27 million particles of PF-06882961 and OWL833 complex respectively) were used to generate the initial 3D model by SGD algorithm for PF-06882961 complex (volume of PF-06882961 complex was used as initial model for OWL833 complex), and subsequently applied to 3D classification. Particles from the best-looking class (1.21 million and 1.72 million particles of PF-06882961 and OWL833 complex, respectively) were subjected to Bayesian particle polishing, CTF refinement and cycles of 3D auto-refinement in RELION (Zivanov et al., 2018). The refined particles were subjected to further 3D classification with fine angular sampling. 683K and 934K particles of PF-06882961- and OWL833-GLP-1R-Gs complexes, respectively, were chosen to generate a final map using 3D auto-refinement and sharpened with a B-factor of −59Å^2^ and −25Å^2^, respectively. Local resolution was determined using RELION with half-reconstructions as input maps. The map density of detergent micelle and Gαs helical domain (AHD) was masked out during the consensus map reconstruction for clarity. The agonist-GLP-1R or ECD of each particle was further refined through receptor/ECD-focused 3D auto-refinement with a loose mask in RELION.

To improve the resolution of AHD, the refined particle stack (636K, 732K and 934K particles of GLP1, PF-06882961 and OWL833 complexes, respectively) was subjected to 3D classification without alignment into 5 classes using a broad mask of the AHD. Each particle class (52K, 74K, 152K, 255K or 102K of GLP1 complex; 115K, 116K, 127K, 158K or 216K of PF-06882961 complex; 152K, 181K, 197K, 200K or 203K of OWL833 complex) was 3D auto-refined, and the best resolved classes (152K, 116K and 181K particles of GLP1, PF-06882961 and OWL833 complexes, respectively) were further refined and post processed with a G protein mask to generate a final map of the AHD.

### Atomic model refinement

*GLP-1-GLP-1R-Gs complex:* The sequence corrected model of ExP5-GLP-1R-Gs (PDB: 6B3J) used as the initial template and fit in the cryo-EM density maps in UCSF Chimera (v1.14) (Pettersen et al., 2004), followed by molecular dynamics flexible fitting (MDFF) simulation with nanoscale molecular dynamics (NAMD) (Chan et al., 2012). The fitted model was further refined by rounds of manual model building in COOT (Emsley et al., 2010) and real space refinement, as implemented in the PHENIX software package (Adams et al., 2010). The ECD and ECLs were modelled manually without ambiguity based on the ECD-focused map. Density of ECD linker (K130^ECD^ −S135^ECD^) and ICL3 (L339^ICL3^-T343^ICL3^) was not observed and these residues were omitted from the final model.

*PF-06882961 or OWL833-GLP-1R-Gs complex:* Geometry restraint information of PF-06882961 and OWL833 was generated using the module ‘electronic Ligand Builder and Optimisation Workbench (eLBOW)’ in PHENIX (Moriarty et al., 2009). The model of GLP-1-GLP-1R-Gs or PF-06882961-GLP-1R-Gs was used as initial template for PF-06882961 or OWL833-GLP-1R-Gs complex modelling, respectively. The template was fit into the cryo-EM density map with the MDFF routine in NAMD (Chan et al., 2012). The fitted models were further refined by manual model building in COOT and real space refinement in PHENIX based on the consensus maps and receptor-focused maps (Adams et al., 2010; Emsley et al., 2010). The model of ECD in the OWL833-complex was further optimized according to the ECD-focused map. Cryo-EM density for residues between S129^ECD^ and R134^ECD^ in the ECD linker region of both small molecule complexes were discontinuous and these sequences were omitted from the final model. The ECL3 was less resolved in OWL833-complex and residues from A375^ECL3^-G377^ECL3^ were omitted.

The AHD of the three complexes was modelled based on their AHD refined maps using the AHD of β2 adrenergic receptor-Gs complex (PDB: 3SN6) as the initial template. Water molecules were modelled manually into clear point density for all the three structures and further refined in Coot and PHENIX (Adams et al., 2010; Emsley et al., 2010). The final models were subjected to global refinement and comprehensive validation implemented in PHENIX. The cryo-EM data collection, refinement and validation statistics for all complexes are reported in Supplementary Table 1.

### Model residue interaction analysis

Interactions in the PDB files between the bound agonist and the receptor were analyzed using the “Dimplot” module within the Ligplot+ program (v2.2) (Laskowski and Swindells, 2011). Hydrogen bonds were additionally analysed using the UCSF Chimera package (Pettersen et al., 2004), with relaxed distance and angle criteria (0.4 Å and 20 degree tolerance, respectively).

### Cryo EM dynamics analysis

3D variability analysis implemented in cryoSPARC (v2.9) (Punjani et al., 2017) was performed to understand and visualize the dynamics in GLP-1R complexes, as previously described for analysis of the dynamics of adrenomedullin receptors (Liang et al., 2020a). The final polished particle stack including GLP-1, PF-06882961 and OWL833-GLP-1R-Gs complexes from RELION consensus refinement were imported into the cryoSPARC environment. 2D classification and selection were initially performed to ensure only highly resolved particles were further analysed. These particles were then subjected to 3D refinement, using a low pass filtered RELION consensus map as an initial model and a generous mask created in cryoSPARC as default. 3D variability of these GLP-1R complexes was analysed in 3 motions and the 20 volume frame data in each motion was generated in cryoSPARC (Punjani et al., 2017). Output files were visualized in UCSF ChimeraX volume series and captured as movies (Goddard et al., 2018).

### Stable cell lines generation

The wild-type (WT) and mutant GLP-1R constructs containing designed signal alanine mutation were integrated into CHOFlpIn cells using FlpIn Gateway technology system (Invitrogen). Stable CHOFlpIn expression cell lines were selected using 600 μg/ml hygromyocin B, and maintained in DMEM supplemented with 5% (V/V) FBS (Invitrogen) at 37°C in 5% CO_2_.

### cAMP accumulation assay – stable cell lines

HEK293A WT GLP-1R, CHOFlpIn WT GLP-1R or CHOFlpIn mutant GLP-1R cells were seeded at a density of 30,000 cells per well into a 96-well plate and incubated overnight at 37 °C in 5% CO_2_. cAMP detection was using a Lance cAMP kit (PerkinElmer Life and Analytical Sciences), performed as previously described (Hager et al., 2017). Growth media was replaced with stimulation buffer [phenol-free DMEM containing 0.1% (w/v) ovalbumin and 1 mM 3-isobutyl-1-methylxanthine] and incubated for 1 h at 37 °C in 5% CO_2_. Cells were stimulated with increasing concentrations of ligand, 100 μM forskolin or vehicle, and incubated for 30 min at 37°C in 5% CO_2_. The reaction was terminated by rapid removal of the ligand-containing buffer and addition of 50 μL of ice-cold 100% ethanol. After ethanol evaporation, 75 μL of lysis buffer [0.1% (w/v) BSA, 0.3% (v/v) Tween 20, and 5 mM HEPES, pH 7.4] was added, and 10 μL of lysate was transferred to a 384-well OptiPlate (PerkinElmer Life and Analytical Sciences). 5 μL of 1/100 dilution of the Alexa Fluor® 647-anti cAMP antibody solution in the Detection Buffer (PerkinElmer Life and Analytical Sciences) and 10 μL of 1/5000 dilution of Eu-W8044 labeled streptavidin and Biotin-cAMP in the Detection Buffer (PerkinElmer Life and Analytical Sciences) were added in reduced lighting conditions. Plates were incubated at room temperature for 2h before measurement of the fluorescence using an EnVision Multimode Plate (Reader PerkinElmer Life and Analytical Sciences). All values were converted to cAMP concentration using cAMP standard curve performed parallel and data were subsequently normalized to the response of 100μM forskolin in each cell line, and then normalized to the WT for each agonist.

### Calcium mobilisation

HEK293A cells stably expressing the hGLP-1R were seeded at a density of 30,000 cells/well into poly-D-Lysine coated 96-well culture plates and incubated overnight at 37°C in 5% CO_2_. Ca^2+^ mobilisation was assessed as previously described (Koole et al., 2010). Briefly, media was changed and cells were incubated with fluo8 for 1 h. Ligand-mediated intracellular calcium mobilisation was determined using a FlexStation (Molecular Devices, USA), with excitation wavelength at 485 nm and emission wavelength at 525 nm every 1.5 sec for 2 minutes, including a 15-second baseline with addition of agonists at increasing concentrations.

### pERK1/2 assays

HEK293 cells stably expressing hGLP-1R were seeded at a density of 30,000 cells/well into Poly-D-Lysine coated 96-well culture plates and incubated overnight at 37°C in 5% CO_2_. Receptor mediated pERK1/2 was determined using the AlphaScreen ERK1/2 SureFire protocol as previously described previously (Wootten et al., 2013a), with increasing concentrations of the compound. Time course experiments were performed using a single ligand concentration to determine the peak response time point (3min) for the subsequent concentration response curves as previously described.

### beta arrestin1/2 assays

HEK 293A Cells were transiently transfected with GLP-1R-Rluc8 and rat β-arrestin1-Venus at a ratio of 1:4 using a standard PEI protocol (6:1 ratio PEI:DNA) and seeded in to poly-D-Lysine coated white 96-well culture plates at a density of 30,000 cells/well. Plates were incubated at 37°C in 5% CO_2_. 48 h post transfection, β-arrestin recruitment was measured in assay buffer (Phenol red-free DMEM, 0.1% (w/v) ovalbumin) as previously described (Savage et al., 2013). The concentration response curves were plotted using the total area under the curve during the time of measurement post ligand addition.

### FYVE trafficking

HEK293A cells were transfected with GLP-1R-Rluc8 and early endosomal marker FYVE tagged with rGFP at 1:4 ratio using a standard PEI protocol (6:1 ratio PEI:DNA) and seeded at a density of 30,000 cells/well in poly-D-Lysine coated white 96-well plate. Following 48 h incubation at 37°C in 5% CO_2_, ligand-mediated receptor trafficking was measured in the presence of 2uM prolume purple substrate in assay buffer (Phenol red-free DMEM, 0.1% (w/v) ovalbumin) as described previously. Plates were read using the PHERAstar FS plate reader (BMG Labtech, Ortenberg, Germany) two wavelengths (emission 1: 410-80nm and emission 2: 515-30nm) at 2 min intervals for 17 min, including a 2-cycle baseline reading preceding compound addition. The concentration response curves were then plotted using the total area under the curve during the time of measurement post ligand addition.

### G_s_ conformational change

HEK293AΔS/Q/12/13 cells stably expressing the GLP-1R (tested and confirmed to be free from mycoplasma) were transfected with a 1:1:1 ratio of Gγ_2_:venus–Gα_s_:nanoluc–Gβ_1_ using a standard PEI protocol (6:1 ratio PEI:DNA). Cells were incubated overnight at 37°C in 5% CO_2_. Collection and preparation of cell plasma membranes and assessment of G protein conformational change was performed using a previously established method (Furness et al., 2016). Briefly, 5 µg per well of cell membrane was incubated with furimazine (1:1,000 dilution from stock) in assay buffer (1× HBSS, 10 mM HEPES, 0.1% (w/v) ovalbumin, 1× P8340 protease inhibitor cocktail, 1 mM DTT and 0.1 mM PMSF, pH 7.4). The GLP-1R-induced BRET signal between Gα_s_ and Gγ was measured at 30°C using a PHERAstar (BMG LabTech). Baseline BRET measurements were taken for 2 min before addition of vehicle or ligand. BRET was measured at 15 s intervals for a further 10 min. All assays were performed in a final volume of 100 μl. The concentration response curves were then plotted using the total area under the curve during the time of measurement post ligand addition.

### NanoBRET kinetic binding assays

HEK293A cells were transiently transfected with Nluc-hGLP-1R. 48 h post transfection, cells were harvested and plasma membrane was extracted as described previously (Furness et al., 2016). 1 μg per well of cell membrane was incubated with furimazine (1:1,000 dilution from stock) in assay buffer (1× HBSS, 10 mM HEPES, 0.1% (w/v) OVA, 1× P8340 protease inhibitor cocktail, 1 mM DTT and 0.1 mM PMSF, pH 7.4). AF568-GLP-1 was used as the fluorescent ligand in the NanoBRET binding assay. Membranes were pre-incubated with PF 06882961 or OWL-833 for 10 min prior to assessment of BRET signal between Nluc-hGLP-1R and AF568-GLP-1 This was assessed using a PHERAstar (BMG LabTech) at 10 sec intervals (25 °C). A 2 min baseline reading was taken before addition of test compound, AF568-GLP-1 (Kd concentration, 3.16nM, determined previously) was added 10 min later, and the measurement continued for 20 min. Data were corrected for baseline and vehicle responses.

### Species selectivity data

#### Transient transfections

Mammalian expression plasmids encoding either human GLP-1R, mouse GLP-1R, human GLP-1R-W33S or mouse GLP-1R-S33W were purchased from ThermoFisher Scientific. Suspension CHO cells were seeded at 0.2-0.3 mill/ml in shaker flasks 2-3 days prior to transfection and cells were propagated to 1-2 mill/ml at 37°C until the day of transfection. Cells were electroporated separately with the 4 different human and mouse GLP-1R expression plasmids and the resulting transfected cells were further cultivated at 37°C for 24 h. One day post-electroporation, cells were subjected to a temperature drop to 32°C, and cells were further cultivated at 32°C for another 24 h until harvest.

#### cAMP assays

Ligand-induced accumulation of cAMP was quantified using the cAMP dynamic Gs kit (Cisbio). Cells were pelleted and washed once in Dulbecco’s Phosphate Buffered Saline (DPBS, ThermoFisher Scientific). Cells were counted, pelleted and resuspended in assay buffer (Hanks’ balanced salt solution buffer (ThermoFisher Scientific), 20 mM HEPES pH 7.4, 1 mM CaCl_2_, 1 mM MgCl_2_, 0,1% Pluronic F-68) to a cell density of 0.5 million cells/mL. Cells (10 µl) were stimulated with increasing concentrations of GLP-1, OWL-833 or PF-06882961 (10 µl) for 30 min at 37°C. The assay was stopped by cell lysis and the response was measured by adding 20 µl/well detection solution (cAMP conjugate and lysis buffer (Cisbio) + 2.5% anti-cAMP cryptate conjugate (Cisbio) + 2.5% cAMP-d2 conjugate (Cisbio). The plate was incubated in the dark for 60 min at room temperature (∼22 °C) and subsequently read on an EnVision Multilabel Reader (PerkinElmer Life and Analytical Sciences), with excitation at 340 nm and measurements of emission at 615 and 665 nm. The fluorescence resonance energy transfer ratios (665/615 nm) were converted to cAMP concentrations by interpolating values from a cAMP standard curve.

### Data analysis

All pharmacological data was analysed using Prism 8.0. Concentration response data were generated by applying a three-parameter logistic equation to either fixed time measurements or data extracted from temporal traces. To quantify kinetic parameters, a one phase association equation was used. Biased agonism was quantified by applying the operational model of agonism to concentration response data where the equation was revised such that the transduction ratio *τ*/K_A_ was directly estimated (Kenakin et al., 2012). Changes to *τ*/K_A_ were established by comparing compounds to the reference ligand, GLP-1, and reference pathway, cAMP production.

### Graphics

Molecular graphics images were produced using the UCSF Chimera (v1.14) and ChimeraX packages from the Computer Graphics Laboratory, University of California, San Francisco (supported by NIH P41 RR-01081and R01-GM129325) (Goddard et al., 2018; Pettersen et al., 2004)

## DATA AND CODE AVAILABILITY

The atomic coordinates and the cryo-EM density maps generated during this study are available at the protein databank (https://www.rcsb.org) and the electron microscopy databank (https://www.ebi.ac.uk/pdbe/emdb) under accession number 6X18 and EMDB entry ID EMD-21992 for the GLP-1:GLP-1R:G_s_ structure, PDB accession number 6X1A and EMDB entry ID EMD-21994 for the PF-06882961:GLP-1R:G_s_ structure, and PDB accession number 6X19 and EMDB entry ID EMD-21993 for the OWL833:GLP-1R:G_s_ structure.

## SUPPLEMENTAL INFORMATION

**Table S1.** CryoEM and model parameters

**Table S2:** Interactions between the GLP-1R and GLP-1, PF-06882961 or OWL833

**Figure S1. Kinetic profiles of binding, transducer coupling and signalling mediated by GLP-1 and non-peptidic agonists.** *Related to* Figure 1. **A,** Agonist-induced changes in trimeric G_s_ protein conformation measured as a ligand-induced change in BRET between Nluc-G*α*_s_ and Venus-G*γ* in membranes prepared from HEK293 cells expressing the GLP-1R. Rates were calculated using a one phase association model; the rate reported was from an approx. EC_90_ concentration of each agonist. **B,** Kinetics of calcium signalling in live HEK293 cells expressing the GLP-1R. **C,** A time-course for pERK1/2 was performed at a single ligand concentration to determine the peak response. This revealed similar profiles of pERK1/2 kinetics, albeit with different maximal signals. **D-E**, Kinetics of *β*-arrestin-1 (D) and *β*-arrestin-2 recruitment (E). **F,** Kinetics for ligand-induced BRET between GLP-1R-Rluc8 and the early endosomal marker FYVE-Venus. Rates were determined using a one phase association model; the rate reported was from an approx. EC_90_ concentration of each agonist. **G,** Kinetic ligand binding assay in membranes measuring nBRET between Nluc-GLP-1R and the fluorescent probe AF568-GLP-1 in the presence and absence of increasing concentrations of unlabelled agonists. The colour key at the bottom provides information regarding the ligand concentrations used for each ligand in each assay. All experiments were performed in GLP-1R HEK293 cells and with the exception of panel G (n=3), all panels show the mean + S.E.M of 5 independent experiments performed in duplicate or triplicate.

**Figure S2. Purification, cryo-EM data imaging and processing of GLP-1R:G_s_ complexes.** *Related to* Figures 2 and 3. **Top,** GLP-1 bound complex; **Middle**, PF 06882961 bound complex; **Bottom,** OWL-833 bound complex. Within each panel: Top left, size exclusion chromatography profile and Coomassie gel of the purified complex. Top middle, 3-D histogram representation of the Euler angle distribution of all the particles used in the reconstruction overlaid on the density map drawn on the same coordinate axis. Top right; exemplar micrograph and 2D class averages of cryo-EM projections of the complex. Bottom left, Gold standard Fourier shell correlation (FSC) curves for the final consensus maps and map validation from half maps, showing the overall nominal resolution. Bottom middle-right, Local resolution-filtered EM maps (consensus and receptor/ECD focused refinements) displaying local resolution (Å) coloured from highest resolution (dark blue) to lowest resolution (red).

**Figure S3. Atomic models of the ligands, receptors and G*α*_s_ *α*5 helix in the cryo-EM map.** *Related to Figures 2, 3, 6 & S7*. EM density map and the model are shown for GLP-1, PF 06882961, OWL-833 and all seven TM helices, and ECLs of the GLP-1R when bound to each agonist. The α5 helix of the Gα_S_ Ras-like domain is also shown for each complex. For GLP-1R, the consensus map was used, with the exception of those labelled * where the ECD refined map is shown. For PF 06882961- and OWL-833 receptor and ligand density was from the receptor-focused refined maps. For the α5 helix, all density shown is from the consensus cryo-EM maps for the respective complex. ECL3 for OWL-833 was poorly resolved and was not modelled.

**Figure S4. Ligand interactions with the GLP-1R.** *Related to* Figure 3. **A,** Overlay of 5VAI with the 2.1 Å GLP-1 bound GLP-1R structure reveals differences within the extracellular region between the two structures. These reveal differences in modelled interactions between GLP-1 and residues within TM1, ECL2/TM5 and TM6/ECL3/TM7. **B,** Top, interactions between the N-terminal portion of GLP-1 (residues 7-17) and the GLP-1R TM bundle as determined by Ligplot+. Middle; Atomic models of the binding sites of the PF-06882961-, GLP-1- and OWL-833-bound GLP-1Rs and their density within the cryo-EM maps. Bottom, interactions between PF 06882961 (left) and OWL-833 (right) with the GLP-1R as determined by Ligplot+. In the Ligplot+ figures, hydrogen bonds are shown as green dotted lines, whereas red dotted lines are hydrophobic contacts. Blue spheres indicate waters that directly interact with ligands.

**Figure S5. Ligand mediated cAMP accumulation in ChoFlpIn cells stably expressing WT or mutant hGLP-1Rs.** *Related to* Figures 3, 4 and 6. Concentration response curves for alanine mutants of the GLP-1R were assessed for GLP-1, PF 06882961 and OWL-833. Residues that either interacted with PF 06882961, OWL-833 or both agonists were selected to assess their importance in ligand-mediated receptor activity. A select set of residues that interact with peptide in the GLP-1 bound structure, but with water in the cryo-EM structures of the GLP-1R bound to PF 06882961 and OWL-833 (right) were also assessed. All data are the means + S.E.M of at least four independent experiments performed in duplicate.

**Figure S6. The GLP-1R binding pocket shape, volume and properties differ in the presence of different agonists.** *Related to Figures 4, 5 and Video S1.* **A,** Slice through of the upper portion of TM bundles bound to different agonists, coloured by hydrophobicity (top) and electrostatic potential (bottom). **B,** Overlay of OWL-833-bound structure with 6ORV (TT-OAD2 bound GLP-1R structure) revealed significant overlap in the ligand binding poses and commonalities in the conformation of TM6 and 7. **C,** Overlay of GLP-1-bound (orange), ExP5 bound (6B3J – yellow), OWL-833-bound (pale blue) and TT-OAD2 bound (6ORV – cornflower blue) GLP-1R structures shows differences in the TM6-ECL3-TM7 conformation.

**Figure S7.** The GLP-1R:G_s_ complexes bound by different agonists share common interactions with G_s_, yet differences in the mobility of the G*α*_s_ AHD. *Related to* Figures 4 and 5 *and Videos S3 and S4.* Top, superimposition of the GLP-1-bound (orange), PF 06882961-bound (pale green) and OWL-833-bound (pale blue) GLP-1R:G_s_ complexes reveal identical backbone conformations within the intracellular portion of the TM bundle, the G protein and the G protein: receptor interface. Rotamer positions of side chains in the lower portion of the receptor bundle were also conserved in the three structures (not shown). **Middle**; Interactions formed between the GLP-1R (orange) and the G protein (*α* - gold (right), *β*-cyan (left)) in the GLP-1 bound GLP-1R structure. Identical interactions were observed in the other two structures, including water (red spheres) mediated interactions. Middle panel shows a cut through of the model with the cryo-EM density showing the G protein:receptor interface. **Bottom,** 3D classification using a mask of the G*α*_s_, including AHD, revealed multiple conformations of this domain relative to the Ras domain. 3 common classes were observed in all three GLP-1R complexes (purple – “open/up”, yellow – “mid” and coral – “closed/down”, however, the % of particles in each class differed depending on the bound ligand; these are shown above the 3D classes. The other two classes differed between complexes and had less well-defined AHDs. The total number of particles used for the 3D classification were 615K, 732K, 933K for GLP-1, PF 06882961 and OWL-833, respectively. While not the most populated, the down conformation (coral) was the only class of sufficient resolution to enable modelling of the AHD and this is shown in panel A.

**Video S1. GLP-1, PF 06882961 and OWL-833 stabilise distinct binding pocket volumes within the GLP-1R**. *Related to* Figures 3-5 *and S6.* Slice through the centre of the GLP-1R bundles bound by different agonists, with a 360° y axis rotation around the centre of each receptor bundle. This highlights distinctions in the volume and shapes of GLP-1R TM domain (TMD) binding cavities of GLP-1, PF 06882961 and OWL-833.

**Video S2. Morphs between the inactive and distinct agonist-bound GLP-1R conformations**. *Related to* Figure 5. Morphs between the inactive GLP-1R (6LN2) and agonist-bound conformations (GLP-1, PF 06882961, OWL-833 and TT-OAD2 (6ORV).

**Video S3. 3D Variability analysis of GLP-1 and non-peptidic agonist bound GLP-1R:G_s_ complexes**. *Related to Figure 2 and 7*. CryoSPARC variability analysis performed on the GLP-1R:G_s_ complexes in the presence of different bound agonists. Transition 1: Principal component 1; Transition 2: Principal component 2; Transition 3; Principal component 3; Transition 4: Static backbone models built into the cryo-EM maps from frame 000 and frame 019 from Principal component 1 of the GLP-1-bound GLP-1R analysis revealed 7.1 - 7.3 Å differences between the maps (when measured from the Cα of D215^ECL1^ and T51^ECD^), highlighting the coordinated movements within ECL1 and the ECD. These were associated with a 4.5 Å movement within the C-terminus of GLP-1 (measured from the Cα of K34^GLP-1^).

**Video S4. 3D Variability analysis reveals differences in the GLP-1 ECD and G_s_ AHD dynamics in the presence of different agonists**. *Related to* Figures 4*, 7 and S7.* Transition 1 shows the cryoSPARC 3D variability analysis performed on each complex. This shows substantial mobility in the G_s_ AHD when the GLP-1R is bound by GLP-1, PF 06882961 and OWL-833, but less mobility in the TT-OAD2 bound complex. In contrast, while the GLP-1R ECD is relatively stable in the PF 06882961, OWL-833 and to a lesser extent GLP-1 bound complexes, this domain is extremely mobile when bound by TT-OAD2. Transition 2 shows morphs between the three common conformations of the G_s_ AHD location for the GLP-1, PF 06882961 and OWL-833 bound GLP-1R:G_s_ complexes determined from 3D classification using a mask of G*α*_s_, including the AHD.

## KEY RESOURCES TABLE

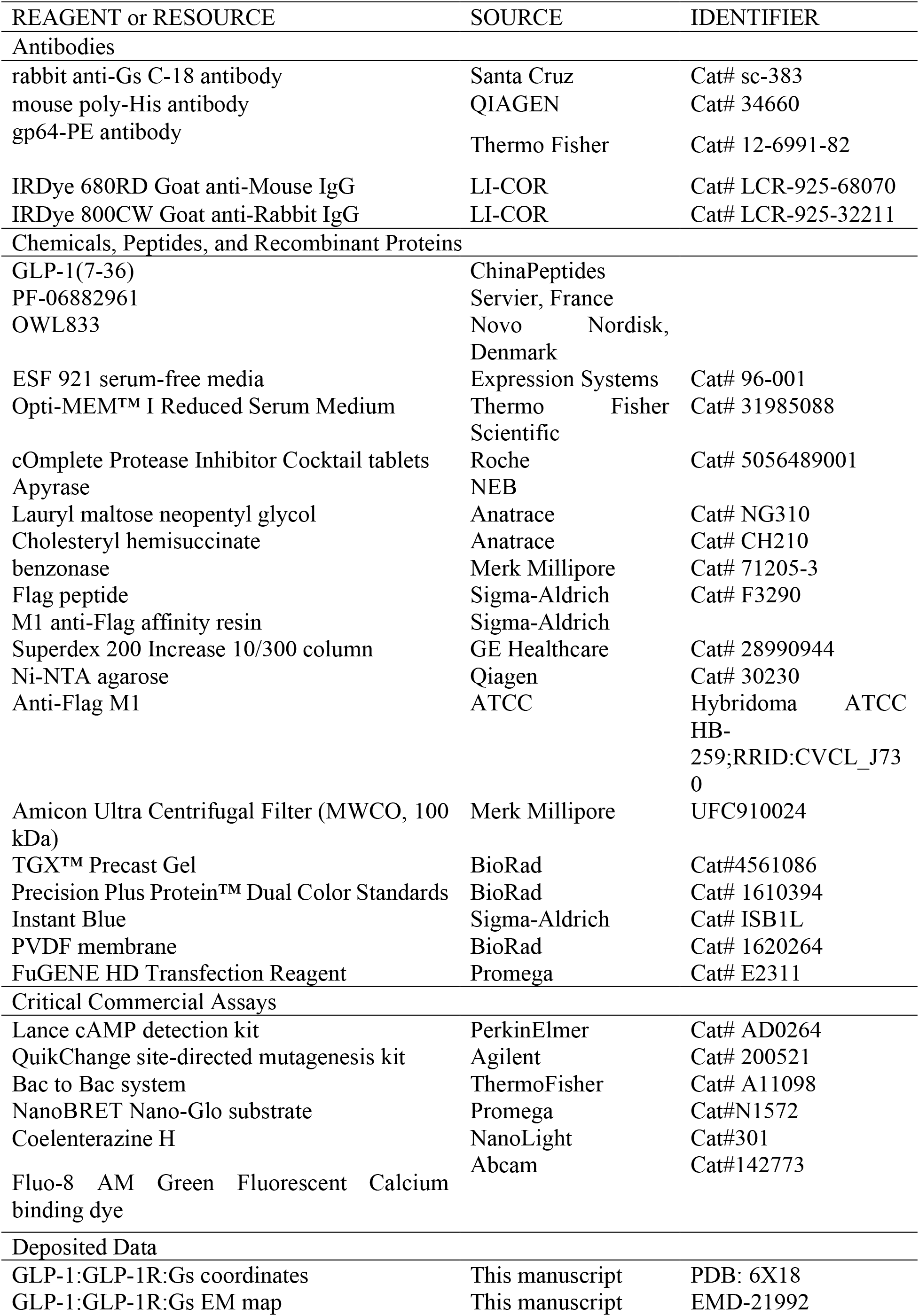
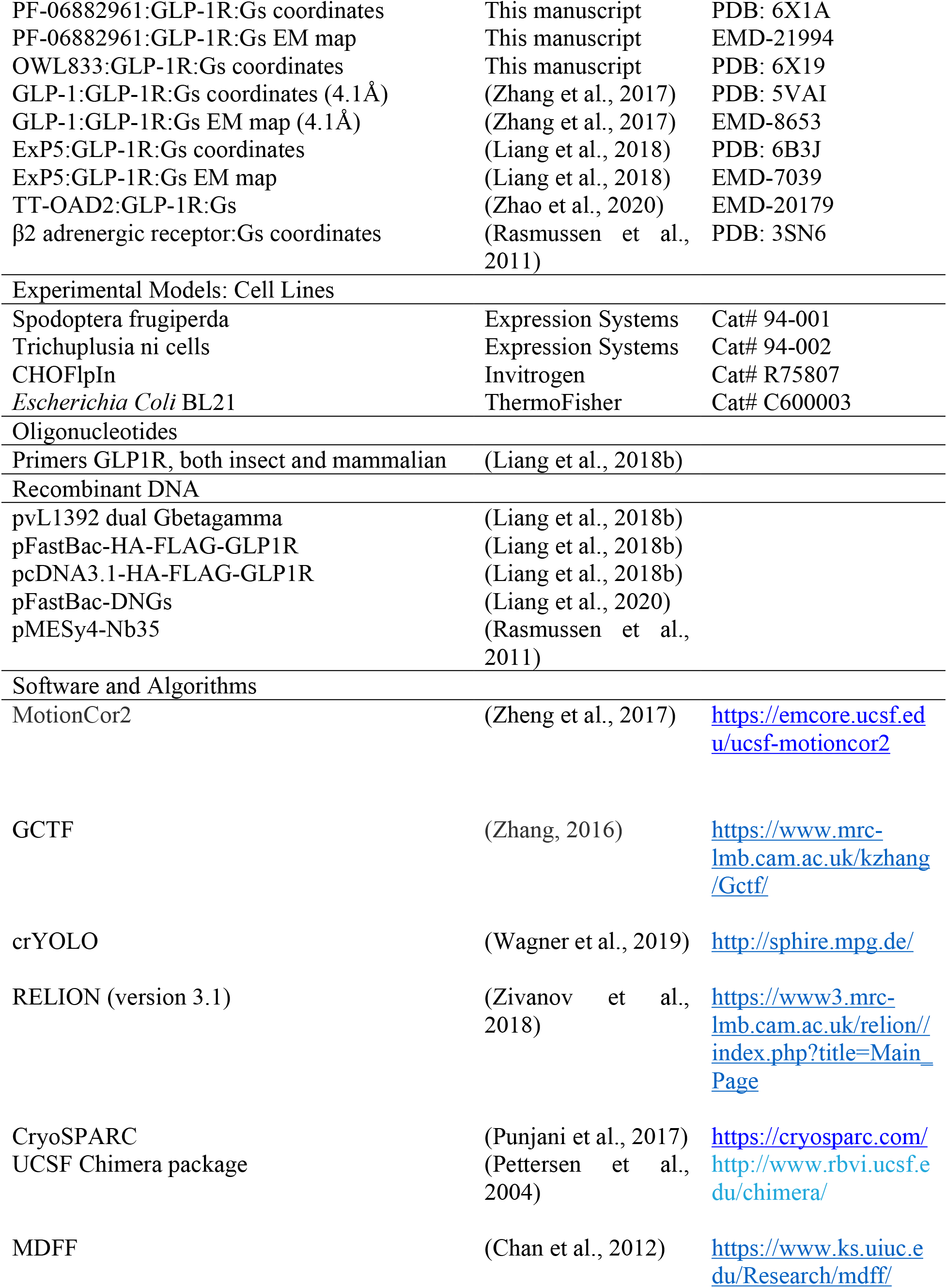
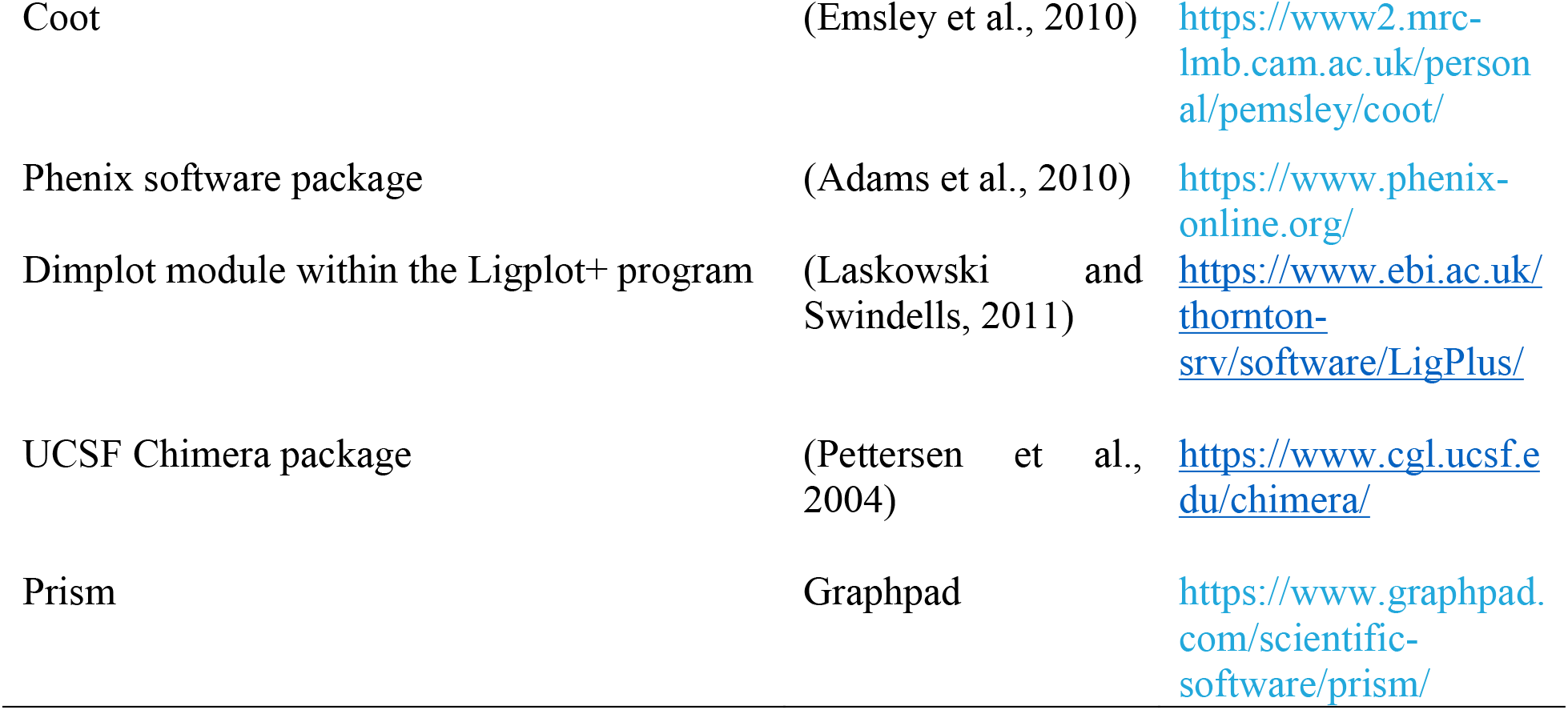

| REAGENT or RESOURCE | SOURCE | IDENTIFIER |
| --- | --- | --- |
| **Antibodies** |  |  |
| rabbit anti-Gs C-18 antibody | Santa Cruz | Cat# sc-383 |
| mouse poly-His antibody | QIAGEN | Cat# 34660 |
| gp64-PE antibody | Thermo Fisher | Cat# 12-6991-82 |
| IRDye 680RD Goat anti-Mouse IgG | LI-COR | Cat# LCR-925-68070 |
| IRDye 800CW Goat anti-Rabbit IgG | LI-COR | Cat# LCR-925-32211 |
| **Chemicals, Peptides, and Recombinant Proteins** |  |  |
| GLP-1(7-36) | ChinaPeptides |  |
| PF-06882961 | Servier, France |  |
| OWL833 | Novo Nordisk, Denmark |  |
| ESF 921 serum-free media | Expression Systems | Cat# 96-001 |
| Opti-MEM™ I Reduced Serum Medium | Thermo Fisher Scientific | Cat# 31985088 |
| cOmplete Protease Inhibitor Cocktail tablets | Roche | Cat# 5056489001 |
| Apyrase | NEB |  |
| Lauryl maltose neopentyl glycol | Anatrace | Cat# NG310 |
| Cholesteryl hemisuccinate | Anatrace | Cat# CH210 |
| benzonase | Merk Millipore | Cat# 71205-3 |
| Flag peptide | Sigma-Aldrich | Cat# F3290 |
| M1 anti-Flag affinity resin | Sigma-Aldrich |  |
| Superdex 200 Increase 10/300 column | GE Healthcare | Cat# 28990944 |
| Ni-NTA agarose | Qiagen | Cat# 30230 |
| Anti-Flag M1 | ATCC | Hybridoma ATCC HB-259;RRID:CVCL_J730 |
| Amicon Ultra Centrifugal Filter (MWCO, 100 kDa) | Merk Millipore | UFC910024 |
| TGX™ Precast Gel | BioRad | Cat#4561086 |
| Precision Plus Protein™ Dual Color Standards | BioRad | Cat# 1610394 |
| Instant Blue | Sigma-Aldrich | Cat# ISB1L |
| PVDF membrane | BioRad | Cat# 1620264 |
| FuGENE HD Transfection Reagent | Promega | Cat# E2311 |
| **Critical Commercial Assays** |  |  |
| Lance cAMP detection kit | PerkinElmer | Cat# AD0264 |
| QuikChange site-directed mutagenesis kit | Agilent | Cat# 200521 |
| Bac to Bac system | ThermoFisher | Cat# A11098 |
| NanoBRET Nano-Glo substrate | Promega | Cat#N1572 |
| Coelenterazine H | NanoLight | Cat#301 |
| Fluo-8 AM Green Fluorescent Calcium binding dye | Abcam | Cat#142773 |
| **Deposited Data** |  |  |
| GLP-1:GLP-1R:Gs coordinates | This manuscript | PDB: 6X18 |
| GLP-1:GLP-1R:Gs EM map | This manuscript | EMD-21992 |
| PF-06882961:GLP-1R:Gs coordinates | This manuscript | PDB: 6X1A |
| PF-06882961:GLP-1R:Gs EM map | This manuscript | EMD-21994 |
| OWL833:GLP-1R:Gs coordinates | This manuscript | PDB: 6X19 |
| GLP-1:GLP-1R:Gs coordinates (4.1Å) | (Zhang et al., 2017) | PDB: 5VAI |
| GLP-1:GLP-1R:Gs EM map (4.1Å) | (Zhang et al., 2017) | EMD-8653 |
| ExP5:GLP-1R:Gs coordinates | (Liang et al., 2018) | PDB: 6B3J |
| ExP5:GLP-1R:Gs EM map | (Liang et al., 2018) | EMD-7039 |
| TT-OAD2:GLP-1R:Gs | (Zhao et al., 2020) | EMD-20179 |
| β2 adrenergic receptor:Gs coordinates | (Rasmussen et al., 2011) | PDB: 3SN6 |
| Experimental Models: Cell Lines |  |  |
| Spodoptera frugiperda | Expression Systems | Cat# 94-001 |
| Trichoplusia ni cells | Expression Systems | Cat# 94-002 |
| CHOFlpIn | Invitrogen | Cat# R75807 |
| *Escherichia Coli* BL21 | ThermoFisher | Cat# C600003 |
| Oligonucleotides |  |  |
| Primers GLP1R, both insect and mammalian | (Liang et al., 2018b) |  |
| Recombinant DNA |  |  |
| pVL1392 dual Gbetagamma | (Liang et al., 2018b) |  |
| pFastBac-HA-FLAG-GLP1R | (Liang et al., 2018b) |  |
| pcDNA3.1-HA-FLAG-GLP1R | (Liang et al., 2018b) |  |
| pFastBac-DNGs | (Liang et al., 2020) |  |
| pMESy4-Nb35 | (Rasmussen et al., 2011) |  |
| Software and Algorithms |  |  |
| MotionCor2 | (Zheng et al., 2017) | https://emcore.ucsf.edu/ucsf-motioncor2 |
| GCTF | (Zhang, 2016) | https://www.mrc-lmb.cam.ac.uk/kzhang/Gctf/ |
| crYOLO | (Wagner et al., 2019) | http://sphire.mpg.de/ |
| RELION (version 3.1) | (Zivanov et al., 2018) | https://www3.mrc-lmb.cam.ac.uk/relion/index.php?title=Main_Page |
| CryoSPARC | (Punjani et al., 2017) | https://cryosparc.com/ |
| UCSF Chimera package | (Pettersen et al., 2004) | http://www.rbvi.ucsf.edu/chimera/ |
| MDFF | (Chan et al., 2012) | https://www.ks.uiuc.edu/Research/mdff/ |
| Coot | (Emsley et al., 2010) | https://www2.mrc-lmb.cam.ac.uk/personal/pemsley/coot/ |
| Phenix software package | (Adams et al., 2010) | https://www.phenix-online.org/ |
| Dimplot module within the Ligplot+ program | (Laskowski and Swindells, 2011) | https://www.ebi.ac.uk/thornton-srv/software/LigPlus/ |
| UCSF Chimera package | (Pettersen et al., 2004) | https://www.cgl.ucsf.edu/chimera/ |
| Prism | Graphpad | https://www.graphpad.com/scientific-software/prism/ |

