## Supplementary Information for "Differential GLP-1R binding and activation by peptide and non-peptide agonists"

### Supplemental Information

#### Table S1. CryoEM and model parameters

| **Data Collection** | *GLP-1-GLP-1R-Gs* | *PF-06882961-GLP-1R-Gs* | *OWL833-GLP-1R-Gs* |
|---|---|---|---|
| Magnification | 105,000 | 105,000 | 105,000 |
| Voltage (kV) | 300 | 300 | 300 |
| Spot size | 4 | 4 | 5 |
| Electron exposure (e-/Å$^2$) | 65.4 | 60.1 | 54.6 |
| Exposure time (s) | 3 | 3 | 5 |
| Movie frames | 75 | 75 | 71 |
| K3 CDS mode | No | No | Yes |
| Defocus range (μm) | 0.6-1.4 | 0.5-1.3 | 0.7-1.5 |
| Pixel size (Å) | 0.826 | 0.826 | 0.826 |
| Symmetry imposed | C1 | C1 | C1 |
| Final particle imaged (no.) | 635,785 | 683,444 | 934,493 |
| Resolution (Å) | 2.1 (2.2[a], 2.4[b], 2.4[c]) | 2.5 (2.5[a], 2.8[b], 2.2[c]) | 2.1 (2.2[a], 2.3[b], 2.8[c]) |
| FSC threshold | 0.143 | 0.143 | 0.143 |
| **Refinement** | | | |
| Initial model used (PDB code) | 6B3J | 6X18 | 6X1A |
| Map sharpening B factor (Å$^2$) | -44 | -59 | -25 |
| Model composition | | | |
| Non-hydrogen atoms | 10371 | 10140 | 10277 |
| Protein residues | 1289 | 1262 | 1258 |
| ligand | 0 | 1 | 1 |
| RMSDs | | | |

| | | | |
|---|---|---|---|
| Bond length (Å) | 0.007 | 0.006 | 0.007 |
| Bond angles (°) | 1.106 | 0.888 | 1.032 |
| Validation | | | |
| MolProbity score | 1.74 | 2.35 | 1.75 |
| Clashscore | 10.83 | 13.98 | 12.45 |
| Rotamer outliers (%) | 1.45 | 3.79 | 1.01 |
| Ramachandran plot | | | |
| Favoured (%) | 97.71 | 96.15 | 97.26 |
| Allowed (%) | 2.21 | 3.85 | 2.66 |
| Disallowed (%) | 0.08 | 0 | 0.08 |

[a]Resolution (Å) of receptor-focused cryo-EM map

[b]Resolution (Å) of ECD-focused cryo-EM map

[c]Resolution (Å) of complex including the “down” conformation of the $G\alpha_s$ AHD

**Table S2: Interactions between the GLP-1R and GLP-1, PF-06882961 or OWL833**

| GLP-1R Receptor residues | | GLP-1 | PF-06882961 | OWL833 |
|---|---|---|---|---|
| ECD | V30$^{ECD}$ | E21 | | |
| | S31$^{ECD}$ | E21 | Pyridine | Methyl pyrazolopiperidine |
| | L32$^{ECD}$ | E21 (H bond)<br>A24<br>F28 | Fluorobenzyl group<br>Ethoxy linker | |
| | W33$^{ECD}$ | | 4-cyano-2-fluorobenzyl group (π-π interaction)<br>Pyridine (π-π interaction) | Indole (π-π interaction)<br>Dimethylmorpholine |
| | E34$^{ECD}$ | | | Methyl pyrazolopiperidine |
| | T35$^{ECD}$ | F28<br>I29 | | |
| | V36$^{ECD}$ | | 4-cyano-2-fluorobenzyl group | |
| | Q37$^{ECD}$ | | 4-cyano group | |
| | W39$^{ECD}$ | F28 (π-π interaction)<br>L32<br>R36 (π-π interaction) | | |
| | E68$^{ECD}$ | L32<br>R36 | | |
| | Y69$^{ECD}$ | I29<br>L32<br>V33 | | |
| | P90$^{ECD}$ | I29 | | |
| | W91$^{ECD}$ | I29 | | |
| | R121$^{ECD}$ | V33 (H bond) | | |
| | E128$^{ECD}$ | K26 | | |
| TM1 | P137$^{1.32}$ | Y19 | | Methyl pyrazolopiperidine<br>Dimethyl fluorophenyl group<br>Imidazolone |

| | | | | |
|---|---|---|---|---|
| | E138$^{1.33}$ | Y19 | | Dimethyl fluorophenyl group<br>Methyl fluoroindazole |
| | L141$^{1.36}$ | F12 | Pyridine | Dimethyl fluorophenyl group |
| | L144$^{1.39}$ | F12 | | Dimethyl fluorophenyl group |
| | Y145$^{1.40}$ | | | Dimethyl fluorophenyl group<br>Methyl fluoroindazole (π-π interaction) |
| | Y148$^{1.43}$ | F12 (π-π interaction) | *GLP-1R F385$^{7.40}$ (π-π interaction)* | Dimethyl fluorophenyl group (π-π interaction) |
| | Y152$^{1.47}$ | E9 (H bond) | Carboxylic acid (water mediated) | |
| TM2 | R190$^{2.60}$ | E9 (H bond) | Carboxylic acid (water mediated) | |
| | K197$^{2.67}$ | T13 (H bond) | Benzimidazole (H bond) | Imidazolone<br>Indole |
| | D198$^{2.68}$ | | | Methyl fluoroindazole<br>Imidazolone |
| | L201$^{2.71}$ | V16 | Piperidine | Dimethylmorpholine |
| | K202$^{2.72}$ | | | Methyl fluoroindazole |
| | W203$^{2.73}$ | | Piperidine | |
| | Y205$^{2.75}$ | S17 (H bond)<br>L20<br>E21 (H bond) | | Methyl fluoroindazole (π-π interaction) |
| ECL1 | S206$^{ECL1}$ | | Fluorobenzyl-oxy group | |
| | T207$^{ECL1}$ | | Fluorobenzyl-oxy group | |
| | Q210$^{ECL1}$ | A24<br>E27 | | |

| | | | | |
|---|---|---|---|---|
| | $W214^{ECL1}$ | F28 (π-π interaction)<br>W31 (π-π interaction) | | |
| | $L217^{ECL1}$ | | 4-cyano-2-fluorobenzyl group | |
| | $Y220^{ECL1}$ | | | Dimethylmorpholine (H bond) |
| | $Q221^{ECL1}$ | E21 | 4-cyano group<br>Oxetane group | |
| TM3 | $C226^{3.29}$ | | | Dimethylmorpholine |
| | $V229^{3.32}$ | | | Dimethylmorpholine |
| | $F230^{3.33}$ | T13 | Oxetane group<br>Benzimidazole-6-carboxylic acid (π-π interaction) | Dimethylmorpholine<br>Indole (π-π interaction) |
| | $M233^{3.36}$ | T13 | Benzimidazole | Dimethylmorpholine |
| | $Q234^{3.37}$ | H7 (H bond) | Carboxylic acid (water mediated) | |
| | $V237^{3.40}$ | H7 | | |
| | $Y241^{3.44}$ | A8 (water mediated)<br>E9 (water mediated) | | |
| ECL2 | $C296^{ECL2}$ | | Oxetane group | |
| | $T298^{ECL2}$ | G10 (water mediated)<br>T13<br>S14<br>S17 (H bond) | 4-cyano group | Indole<br>Methylcyclopropyl group<br>Carbonyl group (water mediated) |
| | $R299^{ECL2}$ | S14<br>S17 (H bond)<br>S18<br>E21 (H bond) | Carboxylic acid (water mediated) | |
| | $N300^{ECL2}$ | G10<br>S14 (H bond) | | Methylcyclopropyl group |
| TM5 | $W306^{5.36}$ | H7 (π-π interaction),<br>G10, T11 | | |
| | $I309^{5.39}$ | H7 | | |

| | | | | |
|---|---|---|---|---|
| | R310$^{5.40}$ | H7 (water mediated) | | |
| | I313$^{5.43}$ | H7 | | |
| ECL3 | D372$^{ECL3}$ | T11 | | |
| TM7 | R380$^{7.35}$ | T11<br>D15 (ionic interaction) | Carboxylic acid (ionic interaction) | |
| | F381$^{7.36}$ | | Piperidine<br>Benzimidazole (π-π interaction) | |
| | L384$^{7.39}$ | A8<br>T11<br>D15 | Benzimidazole | |
| | F385$^{7.40}$ | | Piperidine | |
| | E387$^{7.42}$ | A8 | | |
| | L388$^{7.43}$ | A8<br>E9 (water mediated),<br>F12 | | Dimethyl fluorophenyl group |

**Video S4. 3D Variability analysis reveals differences in the GLP-1 ECD and $G_s$ AHD dynamics in the presence of different agonists**. *Related to Figures 4, 7 and S7.* Transition 1 shows the cryoSPARC 3D variability analysis performed on each complex. This shows substantial mobility in the $G_s$ AHD when the GLP-1R is bound by GLP-1, PF 06882961 and OWL-833, but less mobility in the TT-OAD2 bound complex. In contrast, while the GLP-1R ECD is relatively stable in the PF 06882961, OWL-833 and to a lesser extent GLP-1 bound complexes, this domain is extremely mobile when bound by TT-OAD2. Transition 2 shows morphs between the three common conformations of the $G_s$ AHD location for the GLP-1, PF 06882961 and OWL-833 bound GLP-1R:$G_s$ complexes determined from 3D classification using a mask of $G\alpha_s$, including the AHD.

Figure S1

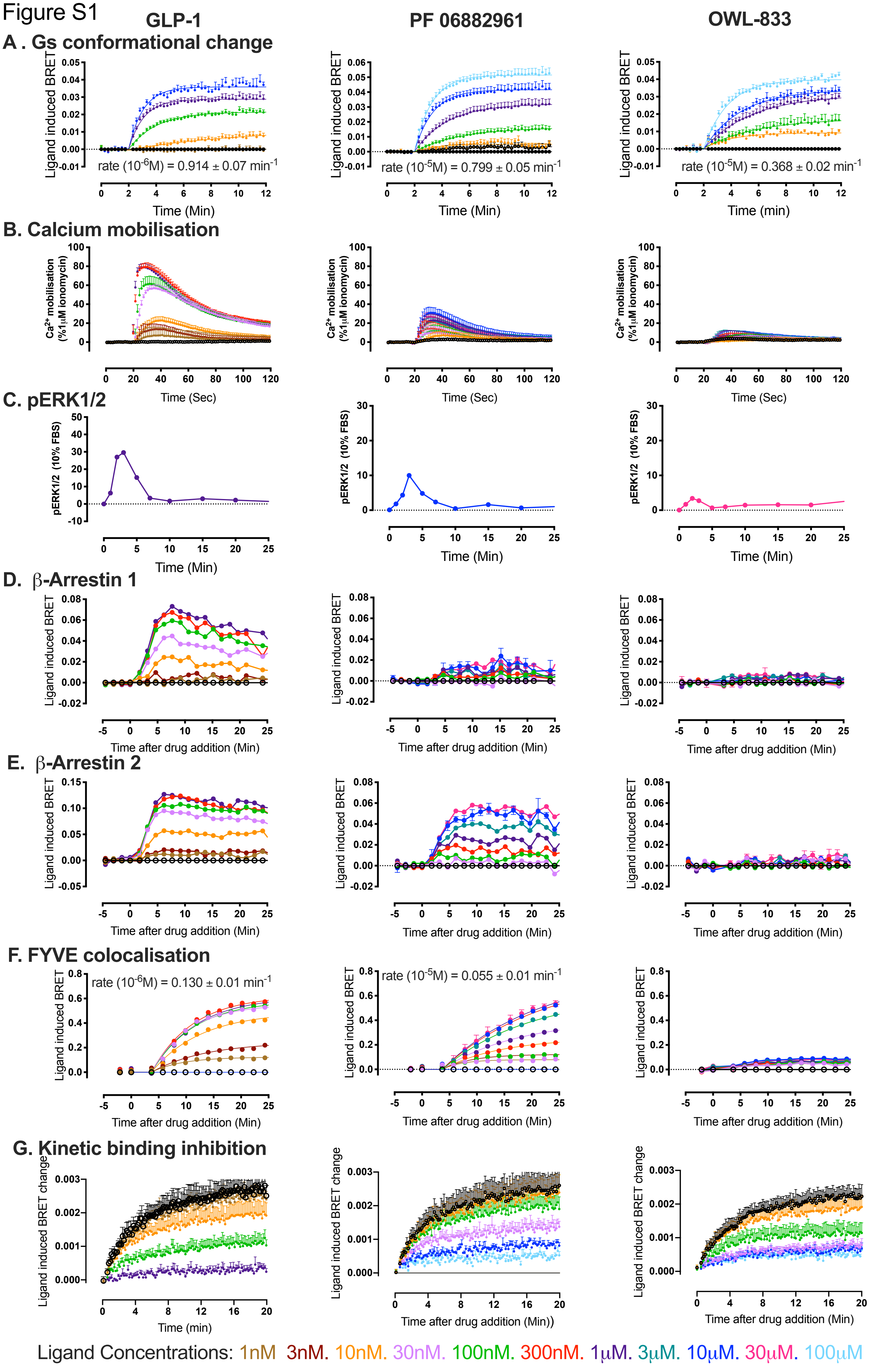
GLP-1
PF 06882961
OWL-833
A . Gs conformational change
Ligand induced BRET
rate (10-6M) = 0.914 ± 0.07 min-1
rate (10-5M) = 0.799 ± 0.05 min-1
rate (10-5M) = 0.368 ± 0.02 min-1
Time (Min)
B. Calcium mobilisation
Ca2+ mobilisation (%1μM ionomycin)
Time (Sec)
C. pERK1/2
pERK1/2 (10% FBS)
Time (Min)
D. β-Arrestin 1
Time after drug addition (Min)
E. β-Arrestin 2
F. FYVE colocalisation
rate (10-6M) = 0.130 ± 0.01 min-1
rate (10-5M) = 0.055 ± 0.01 min-1
G. Kinetic binding inhibition
Ligand induced BRET change
Time (min)
Ligand Concentrations: 1nM 3nM. 10nM. 30nM. 100nM. 300nM. 1μM. 3μM. 10μM. 30μM. 100μM

Figure S2

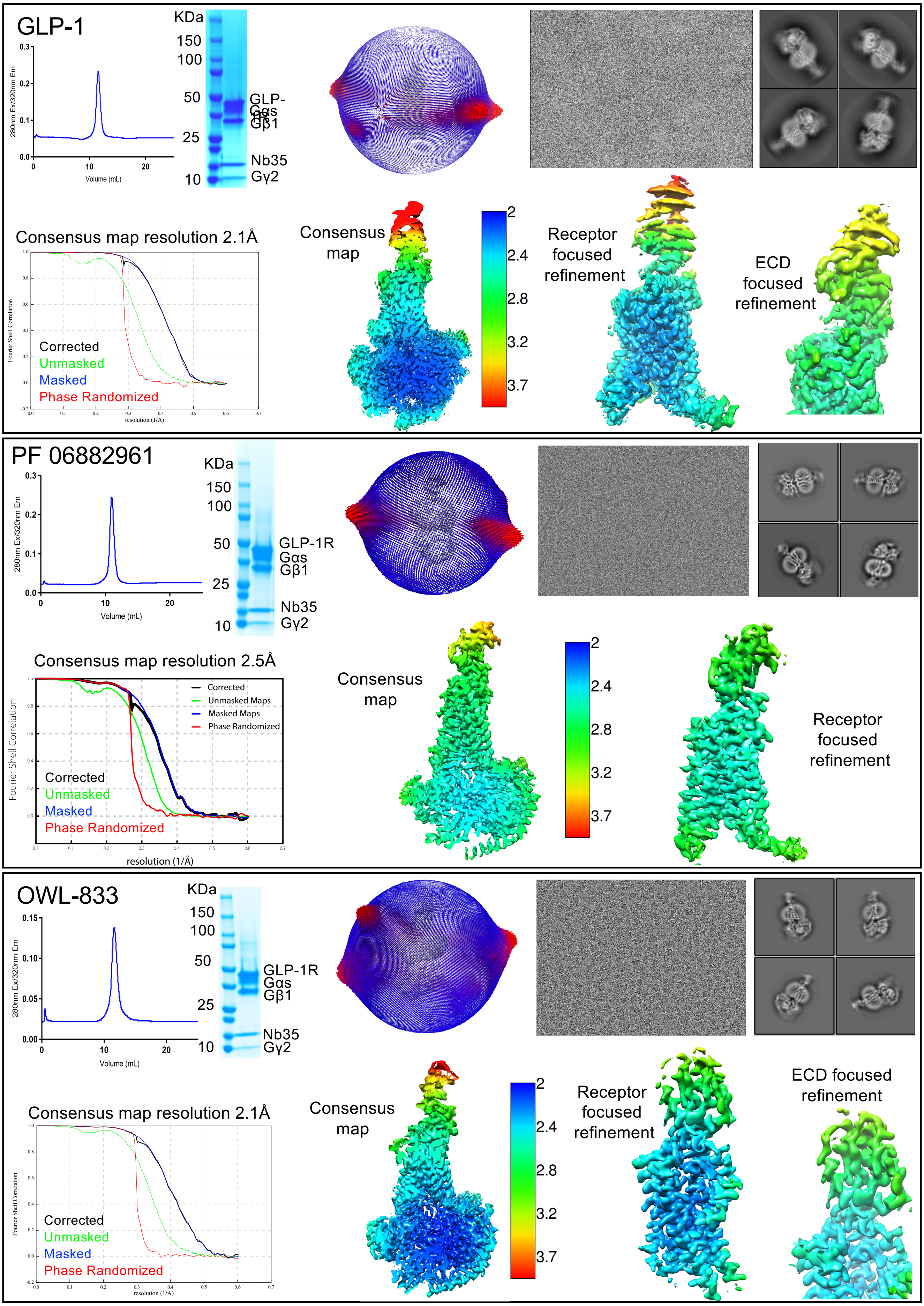
GLP-1
KDa
150
100
50
25
10
GLP-
Gβ1
Nb35
Gγ2
280nm Ex/320nm Em
Volume (mL)
Consensus map resolution 2.1Å
Fourier Shell Correlation
resolution (1/Å)
Corrected
Unmasked
Masked
Phase Randomized
Consensus map
Receptor focused refinement
ECD focused refinement
2
2.4
2.8
3.2
3.7
PF 06882961
KDa
150
100
50
25
10
GLP-1R
Gαs
Gβ1
Nb35
Gγ2
280nm Ex/320nm Em
Volume (mL)
Consensus map resolution 2.5Å
Fourier Shell Correlation
resolution (1/Å)
Corrected
Unmasked Maps
Masked Maps
Phase Randomized
Corrected
Unmasked
Masked
Phase Randomized
Consensus map
Receptor focused refinement
2
2.4
2.8
3.2
3.7
OWL-833
KDa
150
100
50
25
10
GLP-1R
Gαs
Gβ1
Nb35
Gγ2
280nm Ex/320nm Em
Volume (mL)
Consensus map resolution 2.1Å
Fourier Shell Correlation
resolution (1/Å)
Corrected
Unmasked
Masked
Phase Randomized
Consensus map
Receptor focused refinement
ECD focused refinement
2
2.4
2.8
3.2
3.7

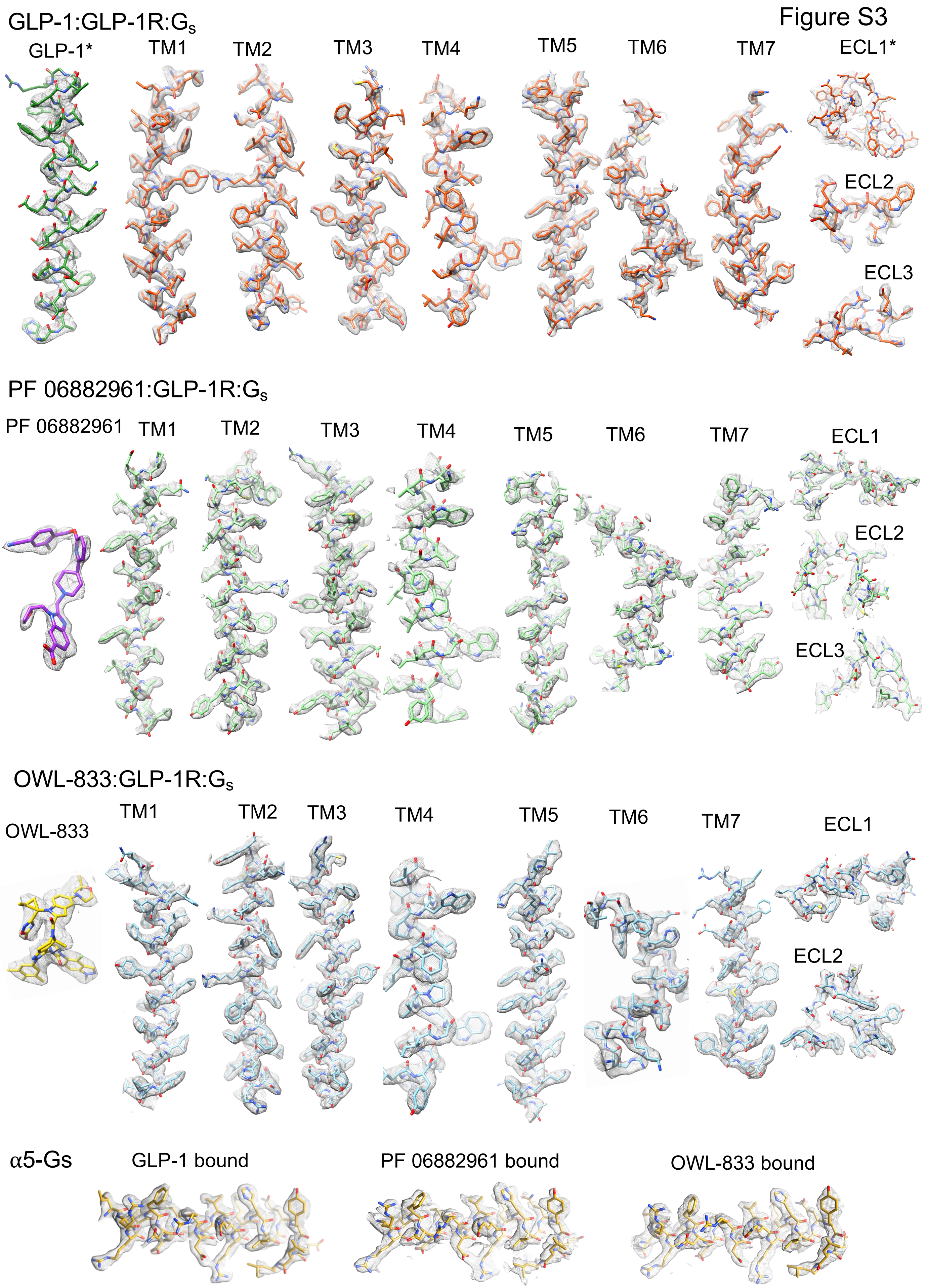

Figure S3
GLP-1:GLP-1R:$G_s$
GLP-1*
TM1
TM2
TM3
TM4
TM5
TM6
TM7
ECL1*
ECL2
ECL3
PF 06882961:GLP-1R:$G_s$
PF 06882961
TM1
TM2
TM3
TM4
TM5
TM6
TM7
ECL1
ECL2
ECL3
OWL-833:GLP-1R:$G_s$
OWL-833
TM1
TM2
TM3
TM4
TM5
TM6
TM7
ECL1
ECL2
α5-Gs
GLP-1 bound
PF 06882961 bound
OWL-833 bound

Figure S4

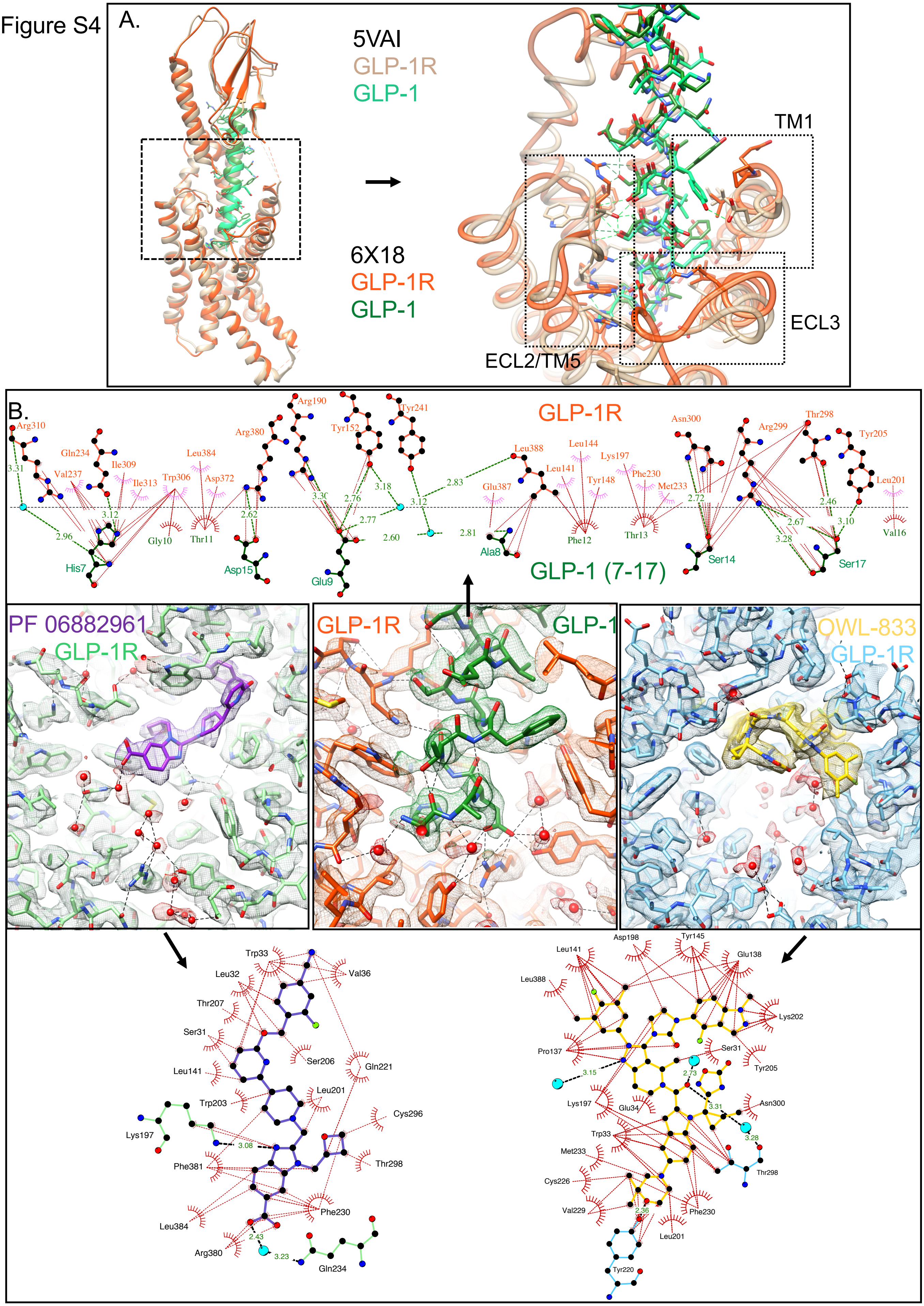
A.
5VAI
GLP-1R
GLP-1
6X18
GLP-1R
GLP-1
TM1
ECL3
ECL2/TM5
B.
GLP-1R
GLP-1 (7-17)
Arg310
Gln234
Val237
Ile309
Ile313
Trp306
Leu384
Asp372
Arg380
Arg190
Tyr152
Tyr241
Leu388
Leu144
Leu141
Glu387
Tyr148
Lys197
Phe230
Met233
Asn300
Arg299
Thr298
Tyr205
Leu201
His7
Gly10
Thr11
Asp15
Glu9
Ala8
Phe12
Thr13
Ser14
Ser17
Val16
3.31
2.96
3.12
2.62
3.30
2.76
3.18
2.77
2.60
3.12
2.83
2.81
2.72
2.67
3.28
2.46
3.10
PF 06882961
GLP-1R
GLP-1R
GLP-1
OWL-833
GLP-1R
Trp33
Leu32
Val36
Thr207
Ser31
Ser206
Gln221
Leu141
Leu201
Trp203
Cys296
Lys197
3.08
Thr298
Phe381
Phe230
Leu384
2.43
Arg380
3.23
Gln234
Asp198
Tyr145
Leu141
Glu138
Leu388
Lys202
Pro137
Ser31
Tyr205
3.15
2.73
3.31
Asn300
Lys197
Glu34
3.28
Trp33
Met233
Thr298
Cys226
Val229
2.36
Phe230
Leu201
Tyr220

Figure S5

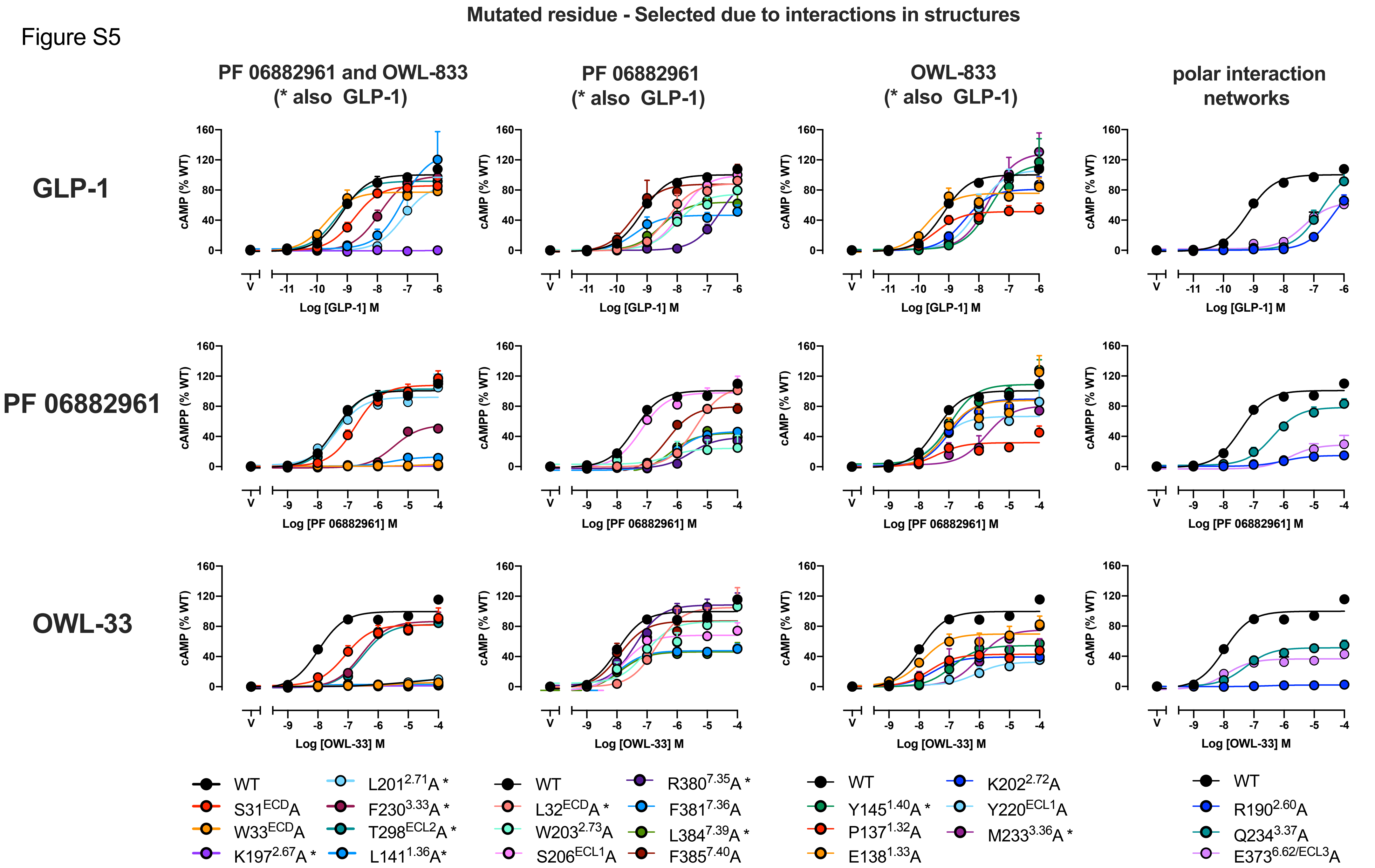

Mutated residue - Selected due to interactions in structures
PF 06882961 and OWL-833 (* also GLP-1)
PF 06882961 (* also GLP-1)
OWL-833 (* also GLP-1)
polar interaction networks
GLP-1
PF 06882961
OWL-33
cAMP (% WT)
cAMPP (% WT)
Log [GLP-1] M
Log [PF 06882961] M
Log [OWL-33] M
WT
S31ECDA
W33ECDA
K1972.67A *
L2012.71A *
F2303.33A *
T298ECL2A *
L1411.36A*
L32ECDA *
W2032.73A
S206ECL1A
R3807.35A *
F3817.36A
L3847.39A *
F3857.40A
Y1451.40A *
P1371.32A
E1381.33A
K2022.72A
Y220ECL1A
M2333.36A *
R1902.60A
Q2343.37A
E3736.62/ECL3A

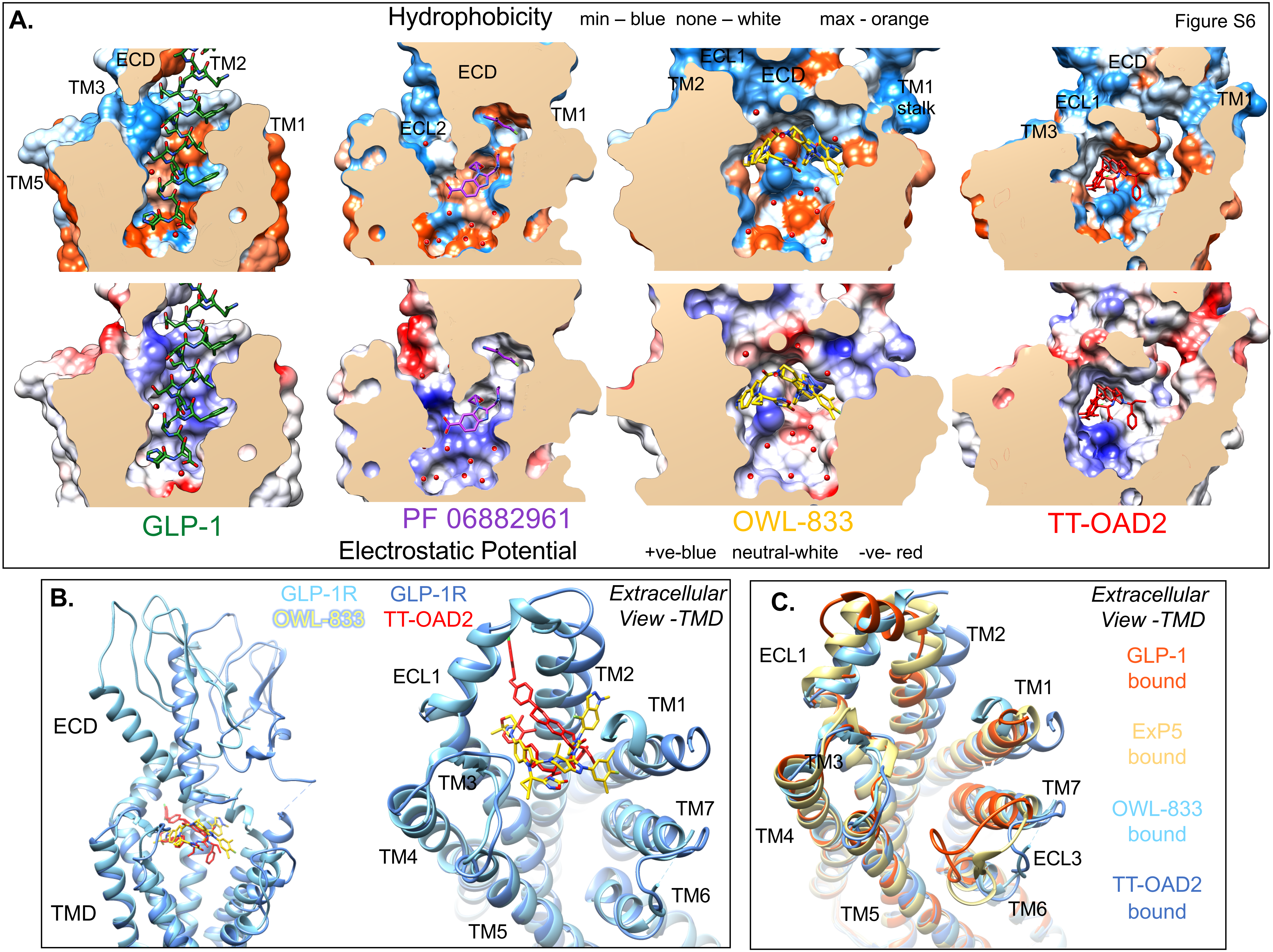
Figure S6
A.
Hydrophobicity min – blue none – white max - orange
ECD
TM2
TM3
TM1
TM5
ECL2
ECL1
stalk
GLP-1
PF 06882961
OWL-833
TT-OAD2
Electrostatic Potential +ve-blue neutral-white -ve- red
B.
GLP-1R
OWL-833
GLP-1R
TT-OAD2
Extracellular View -TMD
ECD
TMD
TM4
TM5
TM6
TM7
C.
GLP-1 bound
ExP5 bound
OWL-833 bound
TT-OAD2 bound
ECL3

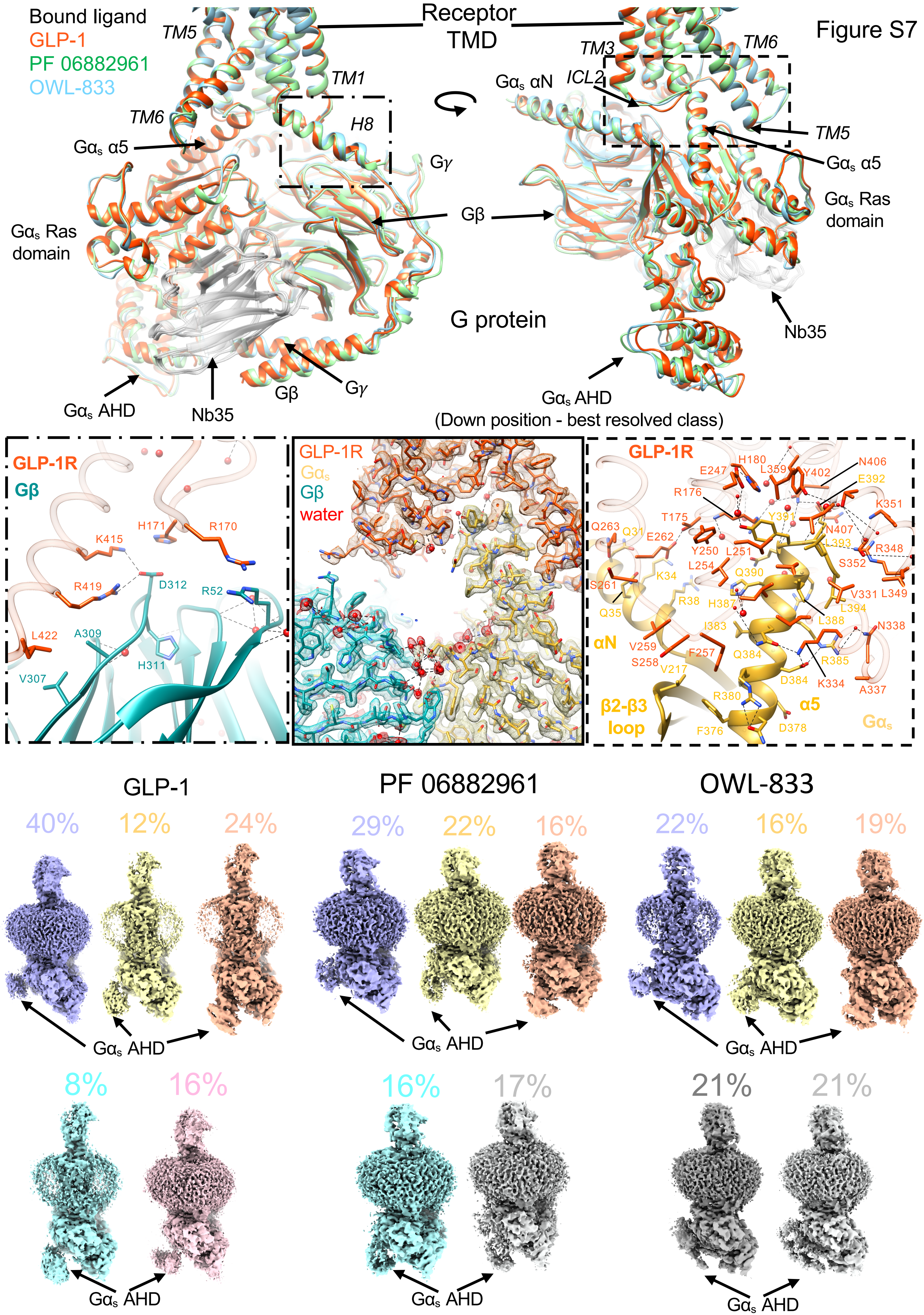

Figure S7
Bound ligand
GLP-1
PF 06882961
OWL-833
Receptor TMD
TM5
TM1
TM6
H8
Gαs α5
Gγ
Gβ
Gαs Ras domain
G protein
Gαs AHD
Nb35
TM3
TM6
Gαs αN
ICL2
TM5
Gαs α5
Gαs Ras domain
Nb35
Gαs AHD
(Down position - best resolved class)
GLP-1R
Gβ
K415
H171
R170
R419
D312
R52
L422
A309
H311
V307
GLP-1R
Gαs
Gβ
water
GLP-1R
H180
N406
E247
L359
Y402
E392
R176
K351
T175
Y391
Q263
Q31
E262
N407
L393
R348
Y250
L251
S352
L254
Q390
K34
S261
R38
H387
V331
L349
Q35
L394
L388
αN
I383
N338
V259
F257
Q384
R385
S258
V217
D384
K334
A337
β2-β3 loop
R380
α5
Gαs
F376
D378
GLP-1
PF 06882961
OWL-833
40%
12%
24%
29%
22%
16%
22%
16%
19%
Gαs AHD
Gαs AHD
Gαs AHD
8%
16%
16%
17%
21%
21%
Gαs AHD
Gαs AHD
Gαs AHD
